# Epromoters bind key stress-related transcription factors to regulate clusters of stress response genes

**DOI:** 10.1101/2024.11.26.625372

**Authors:** Juliette Malfait, Jing Wan, Himanshu Singh, Charbel Souaid, Gaëlle Farah, Cyril Esnault, Sandrine Sarrazin, Michael Sieweke, Salvatore Spicuglia

**Author notes:** Corresponding author: Salvatore Spicuglia. These authors contribute equally (Jing Wan, Himanshu Singh, Charbel Souaid).

## Abstract

Stress insults trigger the rapid and global reprogramming of gene transcription by coordinated recruitment of a limited number of key inducible transcription factors to cis-regulatory elements. Here, we performed a comprehensive analysis of different stress models and observed that co-induced genes are generally located in close genomic proximity. By integrating gene expression and transcription factor binding resources in different stress models, we found an enrichment for clusters whereby only one of the promoters of the cluster recruits the key transcription factors, reminiscent of Epromoters, a type of cis-regulatory elements displaying both promoter and enhancer function. Epromoter-regulated clusters were frequently found irrespectively of the stress or inflammatory response. Predicted Epromoters displayed enhancer activity and regulated clusters of stress-response genes independently of their genomic location. These findings have significant implications for understanding complex gene regulation following the response to acute perturbations.

**Teaser:** When cells face stress, they undergo rapid changes in gene expression, orchestrated by a handful of key transcription factors. But how do these factors coordinate such a complex response? Our study reveals that “Epromoters”—cis-regulatory elements that combine the functions of both promoters and enhancers help organize stress-response genes into tightly regulated clusters. This discovery not only deepens our understanding of gene regulation in the face of stress but also offers exciting implications for studying inflammation and other acute cellular responses.

## Introduction

Cells and organisms are constantly exposed to extrinsic and intrinsic stressors that endanger homeostasis and fitness (*1*). Such stresses can be induced by hypoxia, toxins, mechanical stimuli, elevated temperature or pathogen infections which all compromise cell structure and function by causing macromolecular damage. To survive such stress insults, cells launch survival programs that mitigate characterized by the activation of rapid and transient transcriptional reprogramming (*2*). In contrast to cell differentiation programs, the stress programs are mediated by a limited number of inducible transcription factors (*3, 4*). For instance, heat-shock (HS) response is mediated by the HSF1 and HSF2 TFs, Hypoxia is regulated by the HIF1 complex and TNFα response by the NF-kB complex. Moreover, previous observations have suggested that co-induced stress-response genes are frequently organized in clusters, suggesting the sharing of *cis*-regulatory elements (*5–9*).

Gene transcription in higher eukaryotes relies on a diverse network of regulatory elements, including promoters and enhancers, which traditionally have been seen as distinct types of regulatory elements. Promoters, located near the transcription start site (TSS), initiate local gene transcription, whereas enhancers, positioned farther from the TSS, activate gene expression over larger distances. However, their distinction has become more blurred in recent years notably by the discovery of Epromoters, cis-regulatory sequences that combine architectural and functional characteristics of both promoters and enhancers (*10–19*). Epromoters can regulate distal promoters when assessed in episomal reporter systems or high-throughput reporter assays in different cellular contexts from drosophila to humans (*10, 11, 13, 15, 18, 20*), while their deletion or epigenetic silencing in their natural context results in the loss of expression of distal genes (*14–18*). Moreover, genetic variation within Epromoters can potentially influence the expression of distal genes (*15, 21–24*).

Previous studies have suggested a link between the Epromoter function and the stress responses, particularly the regulation of interferon-response genes (*15, 18, 20, 25*). Using high-throughput reporter assays we previously showed that interferon alpha (IFNα)-induced genes are frequently organized in clusters regulated by a single Epromoter (*18*). Remarkably, within these clusters, Epromoters function as regulatory hubs that selectively recruit transcription factors (TFs) essential for IFNα-driven responses, coordinating gene expression across the cluster. It is likely that this type of regulatory element could be more generally involved in the rapid response of genes to cellular stress. This idea fits well with the notion that co-regulated genes are often located in close genomic proximity of defining inducible transcription factories containing a high concentration of RNA polymerase and key TFs and where efficient transcription can be triggered (*7, 12, 18, 26, 27*). Noteworthy, many of the regulatory elements of inducible genes, such as metallothioneins, histones of early cleavage stages, viral immediate-early genes (from some papovaviruses, cytomegaloviruses, and retroviruses), heat-shock genes and the antiviral interferon genes are located close to, or overlapping with, the promoter region of these genes (*28, 29*). A common characteristic of most of the aforementioned promoters is that they are associated with inducible genes that have to quickly respond to environmental stress.

By analyzing a comprehensive dataset of stress and inflammatory responses from different cellular and environmental cues in mice and humans, we found that induced genes are preferentially found in close genomic proximity. We hypothesized that stress-induced genes are globally organized into clusters, with Epromoters regulating the clustered genes by recruiting key inducible transcription factors in order to obtain a fast and coordinated response to the stress insults. To explore Epromoter function upon different stress insults, we developed a pipeline to identify Epromoters and their potential target genes across various stress conditions, bypassing the need for high-throughput enhancer assays. By retrieving transcriptomic and ChIP-seq data from 51 studies, we identified Epromoter-regulated clusters across 18 distinct stress conditions. Reporter assays and CRISPR-Cas9 genomic deletions confirmed Epromoter activity and demonstrated their regulatory influence on neighboring genes within these clusters at their endogenous loci. Additionally, repositioning an Epromoter within its locus demonstrated its intrinsic enhancer and promoter functions, underscoring the dual role of Epromoters in gene regulation. Thus, Epromoters have a broad role in the induction of co-regulated genes during the mammalian stress response.

## Results

### Stress-response genes are preferentially found in genome proximity

To better understand the regulation of stress-response genes, we selected a series of published stress-response datasets from human and mouse, including gene expression data (RNA-seq, Pro-seq or microarrays) and ChIP-seq for key transcription factors involved in the same stress responses. We compiled data from 28 studies, comprising 32 stress or stimulation conditions for a total of 51 datasets (*6, 18, 30–55*). The datasets cover 18 different stress or stimulation conditions, such as heat shock, serum starvation/response, hypoxia, TNFα, IFNα, IFNγ, and LPS (Table S1A). For example, the “Vihervaara” dataset (*30*) consisted of Pro-seq data and ChIP-seq of transcription factors HSF1 and HSF2 before and after K562 cells were exposed to HS at 42°C for 30 minutes.

Previous studies have suggested that Epromoters might regulate co-induced genes located in genome proximity (*18*). Therefore, we first aimed to determine whether there was a bias in the genomic distribution of stress-induced genes. We calculated the distance between the transcription start sites (TSS) of the closest gene pairs for each set of stress-induced genes (Fig. 1A) and compared their distance distribution to the same number of randomly selected genes (Fig. 1B upper panels; Fig. S1). As a control, we analyzed data from embryonic stem cell (ESC) differentiation into three germ layers (mesoderm, ectoderm, and endoderm) (*56*). We observed that the majority of stress-induced genes were located closer to each other compared to a random distance distribution. For example, HS and TNFα induced genes have a distribution summit of 57.6 kb and 63.8 kb as compared with 136 kb and 119 kb for random genes, respectively (*P* value: 3e^-09^ and 6.7e^-08^; respectively K-S test). However, no significant differences were observed for the distance distribution of induced genes after ESC differentiation in any of the three germ layers (Fig. 1B upper panels; Fig. S1).

**Fig. 1.**
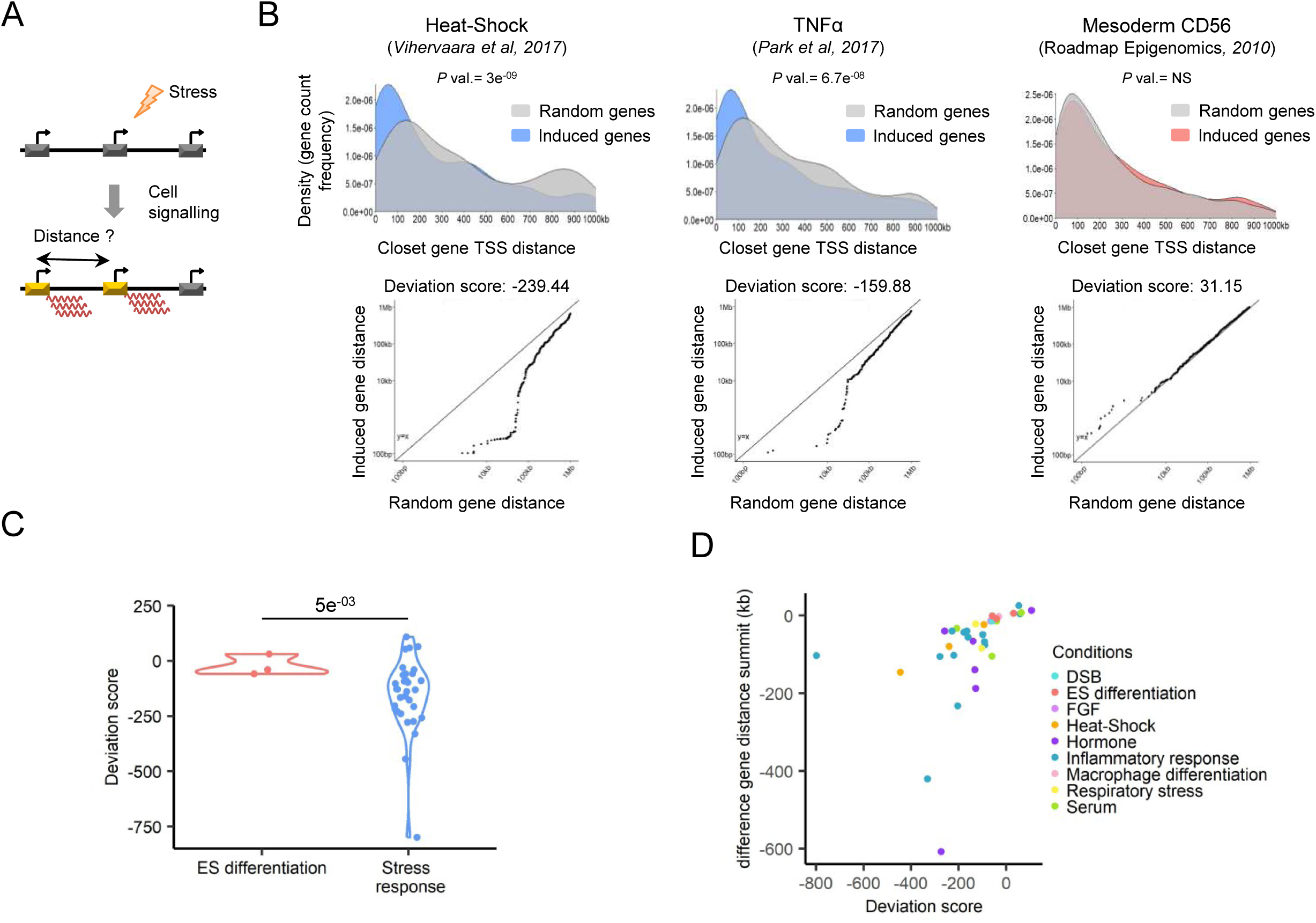
Genomic distance between stress responses induced genes. (A) Approach to assess the distance between two induced genes upon stress. (B) (top panel) Distance distribution of the induced genes (colors) compared to the same numbers of random genes (grey) by dataset (blue: heat shock, and TNFα, red: Mesoderm). *P* values were calculated by Kolmogorov–Smirnov (KS) test. (bottom panel) Deviation scores were calculated between the distance distribution of induced and random genes. (C) Violin plot of the deviation score between datasets from embryonic stem (ES) differentiation (pink) or stress responses datasets (blue). Each point represents a dataset. P value was calculated by a two-sided Student’s t-test. (D) Distribution of each dataset deviation score (x-axis) compared to their difference between the induced and random gene distance summit (y-axis). The dataset is classified into 9 conditions highlighted by different colors (light blue: double-strand breaks (DSB), salmon: ES differentiation, light purple: Fibroblast growth factors (FGF) orange: heat shock, purple: hormone, blue: inflammatory response, pink: macrophage differentiation, yellow: respiratory stress, green: serum.

To assess the distribution biases more precisely, we also calculated a deviation score based on the difference in TSS distances between the random and induced genes (Fig 1B, bottom panels; Fig. S1). The deviation score could be positive or negative depending on whether induced genes are more distant or closer than random genes, respectively. For instance, the HS and TNFα induced genes have a negative deviation score of −239.44 and −159.88, respectively, while the mesoderm-induced genes have a positive deviation score of 31.15. Globally, stress response genes have a significantly lower deviation score than genes induced after ESC differentiation, indicating a closer distance between stress-induced genes (Fig. 1C-D). We noted, however, that 8 out of 32 stress conditions displayed higher or similar distant distribution than ESC-induced genes, highlighting variability in gene organization depending on the type of stress (Fig. 1C-D). Taken together, these results suggest that genes induced by the stress response tend to be located in proximity, suggesting they might be organized into stress-response clusters of co-regulated genes.

### Identification of HS clusters regulated by Epromoters

We previously observed that within IFNα-induced clusters, Epromoters, functionally identified by STARR-seq, preferentially recruited the key TFs STAT1/2 and IRF1/9 (*18*). We wondered whether we could take advantage of this observation to predict Epromoters in different stress or stimulatory conditions. We reasoned that within a cluster of induced genes in response to extracellular or intracellular signaling, the promoter that preferentially recruits the key TFs can be predicted to function as an Epromoter. We developed a pipeline to predict promoters with Epromoter activity upon a stress response (Fig. 2A). First, we identified clusters of induced genes in two ways: either the TSS of induced genes were located less than 100 kb from each other, or they were within the same topologically associating domain (TAD) (Fig. 2A-B). The threshold of 100 kb was defined based on the distance distribution summit between induced genes (range between 50 kb and 90 kb; Fig. 1) as well as previous findings about *cis*-regulatory networks (*18, 31, 57, 58*). Secondly, we assessed whether the stress key TF binding occurred within a [-1; +1] kb window centered on the TSS. We then counted how many promoters within each cluster bound the TF, considering a cluster potentially regulated by an Epromoter if only one promoter was bound by the key TFs (Fig. 2A). We used this prediction to explore the function of Epromoters in the different stress responses mentioned above.

**Fig. 2.**
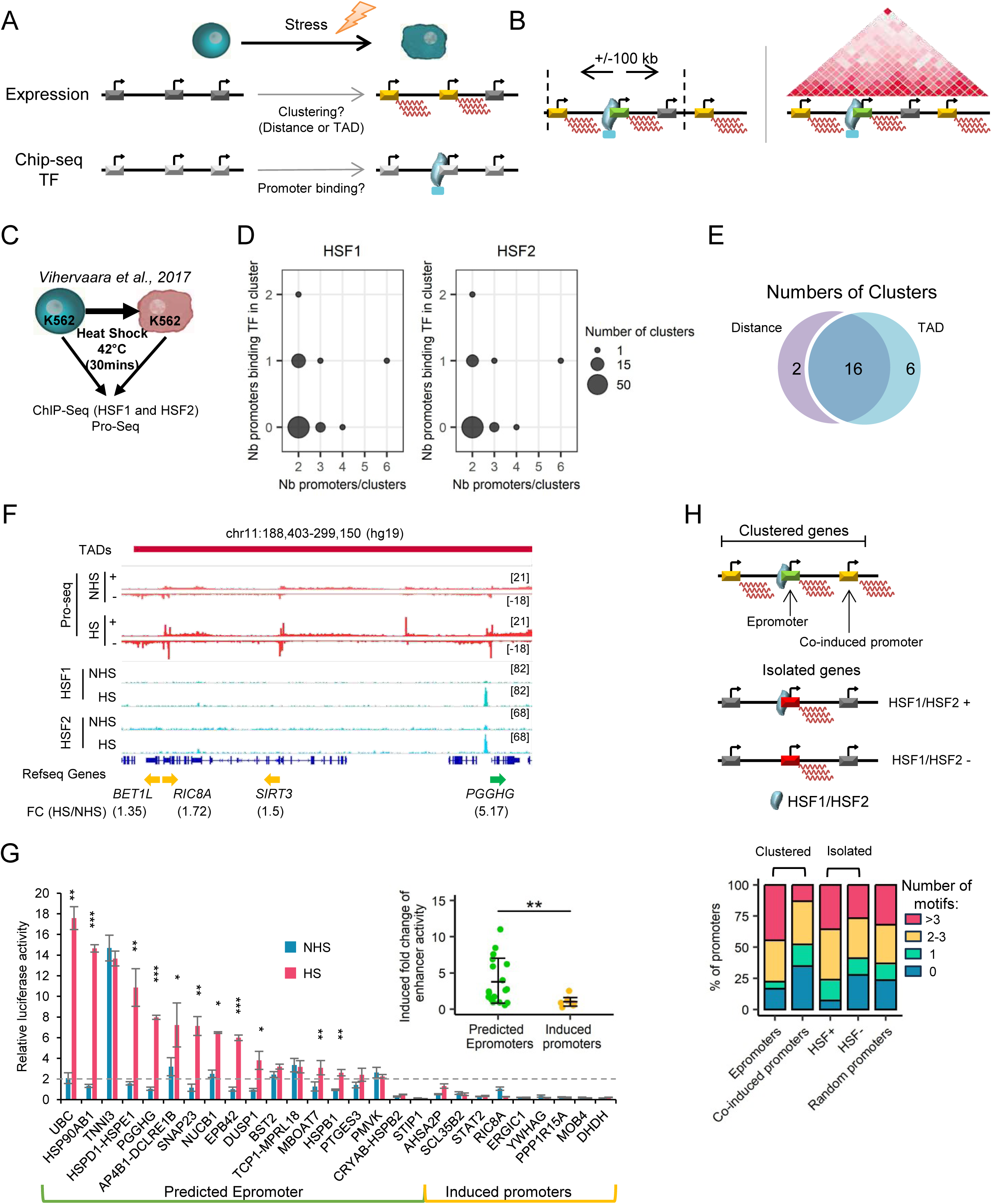
Identification and validation of Epromoter-regulated clusters in the HS stress response. (A) Schematic representation of the pipeline steps to identify Epromoter-regulated gene clusters. (B) Schematic overview of the two clustering methods used in the pipeline. (Left) Clustering of stress-induced genes based on the proximity of their TSS within a 100 kb distance. (Right) Clustering of stress-induced genes based on their localization within the same TAD. (C) Schematic representation of the “Vihervaara” dataset to induce the HS response. (D) (left) Bubble plot showing the number of all the clusters identified by their number of promoters (x-axis) compared to their number of promoters recruiting HSF1 (y-axis) determined by the pipeline. (right) Bubble plot showing the number of all the clusters identified by their number of promoters (x-axis) compared to their number of promoters recruiting HSF2 (y-axis) determined by the pipeline. (E) Venn diagram displaying the overlap of Epromoter-regulated clusters identified after clustering either by the distance between the TSS or their localization within the same TAD in the HS response. (F) Example of the *PGGHG* Epromoter-regulated cluster identified by the pipeline with the two clustering methods in the HS stress response. The genomic tracks show the PRO-seq signal (red) and HSF1 and HSF2 ChIP-seq signal (blue) before or after HS from the “Vihervaara” dataset. The topological associating domains (TAD) from K562 is displayed on the top. The fold-change of induction is indicated below the name and orientation of the genes (Epromoter: green, co-induced genes: yellow). (G) Luciferase assays to quantify the enhancer activity of predicted Epromoter (on the right) and induced promoters clustered with Epromoters (on the left) before (blue) and after (red) HS in the K562 cells. The results were normalized on the pGL4-SV40 promoter plasmid. *P* values were calculated by a paired one-sided Student’s t-test. (insert) Luciferase assay-induced enhancer activity represented by the fold change before/after HS for the predicted Epromoter and the induced promoters. *P* values were calculated by a two-sided Wilcoxon’s test, \*\*\**P* < 0.001, \*\**P* < 0.01, \**P* < 0.1. (H) (top panel) Representation of the classification of the HS-induced genes in 4 categories. (bottom panel) Percent stacked barplot of the numbers of motifs (HSE) per promoter in each category. The number of motifs is divided into 0 motif (blue), 1 motif (cyan), 2 to 3 motifs (yellow), and more than 3 motifs (red).

We compared the effectiveness of the two clustering methods using the “Vihervaara” dataset (*30*) HS in K562 cells (Fig. 2C). Analysis of 778 HS-induced genes using the distance-based method resulted in 68 clusters, harboring 221 genes. We distributed those clusters by the number of promoters per cluster in function of the number of promoters recruiting the HS TFs (here HSF1 and HSF2). We found 18 clusters where only one promoter bound HSF1 and HSF2 (Fig. 2D, Table S2). This suggests that in HS, 18 induced clusters might be regulated by an Epromoter. Using the TAD-based method resulted in 106 clusters, harboring 261 genes, and predicted 22 Epromoter-containing clusters. Comparison between the two methods resulted in 16 common clusters (Fig. 2E). The TAD-based method identified six additional clusters likely due to the lack of threshold between the TSS distances, while the two clusters uniquely identified by the distance-based method might be explained by the overlap of the cluster with the TAD borders (Fig. S2A-B). Nevertheless, the two methods provided consistent identification of gene clusters potentially regulated by Epromoters. An example of an HS cluster potentially regulated by an Epromoter is provided by the *PGGHG* Epromoter-regulated cluster, including three promoters for four induced genes (one promoter is bidirectional) within the same TAD. Among the three promoters, only the *PGGHG* promoter recruited HSF1 and HSF2, suggesting that the HS-induction of *SIRT3*, *RIC8A*, and *BET1L* is mediated by the *PGGHG* Epromoter (Fig. 2F). For this analysis, we used TAD data from K562 cells without HS. Given that TAD data were not always available for the relevant cell lines and genome annotations and considering the potential need for stress-specific topological information, we focused, for the rest of the study, on clusters identified by the distance-based method.

To validate the efficiency of the pipeline in identifying Epromoter-regulated clusters, we tested the enhancer activity of the 18 HS-induced Epromoters from the “Vihervaara” dataset by luciferase reporter assay in K562. As controls, we also tested 9 co-induced promoters. HS induction was confirmed by measuring the promoter activity of the HSPA1A promoter (Fig. S2C). After transfection, cells were incubated for 1 hour at 42°C followed by 2 hours of recovery at 37°C. Out of the 18 predicted Epromoters, 16 exhibited enhancer activity (at least 2-fold induction), among them 11 Epromoters displayed significant induction of enhancer activity upon HS (Fig. 2G). In contrast, none of the 9 co-induced promoters showed enhancer activity in these assays. Overall, compared to the induced promoters, the Epromoters displayed significant HS-induced enhancer activity (Fig. 2G, insert, Fig. S2D). The high proportion of experimentally validated HS-response Epromoters supports the efficiency of our strategy in identifying stress-response Epromoters.

Previous studies by Santiago et al., have shown that Epromoters in the IFNα response possess a higher density of TF binding sites (TFBS) specific to this stress (*18*). We sought to verify whether this trend holds true for the newly identified Epromoters in the HS stress response. Using the JASPAR database, we computed the number of HSF1 or HSF2 binding sites (also called HSEs for Heat shock Elements) *per* HS-induced promoters (Fig. 2H, top panel). To compare the number of motifs between promoters, we divided the induced gene promoters into four categories: the predicted Epromoters, the predicted co-induced promoters (clustered with Epromoters), and all the other induced gene promoters whether they recruit the TFs (HSF+) or not (HSF-). Additionally, we included a random set of promoters as a control for the background of HSE presence. We observed that Epromoters and HSF+ promoters displayed a higher density of HSE motifs compared to co-induced and HSF-promoters, respectively. For instance, 44% of Epromoters harbor 3 or more HSE motifs while this is the case for only 13% of co-induced promoters. Strikingly, we also observed that co-induced promoters harbored fewer HSEs than other HS-induced promoters or random promoters, suggesting their depletion of HSEs. These observations support a model whereby the co-induced promoters are dependent on the binding of HSF1/HSF2 at Epromoters within the same cluster. Furthermore, we assessed the functional relevance of HS-induced genes (Fig. 3E). As expected, we found that Epromoter-genes were enriched in Biological Process GO terms associated with heat shock, such as response to unfolded protein and protein stabilization and folding similar to other induced genes, while co-induced genes displayed no significant enrichment.

**Fig. 3.**
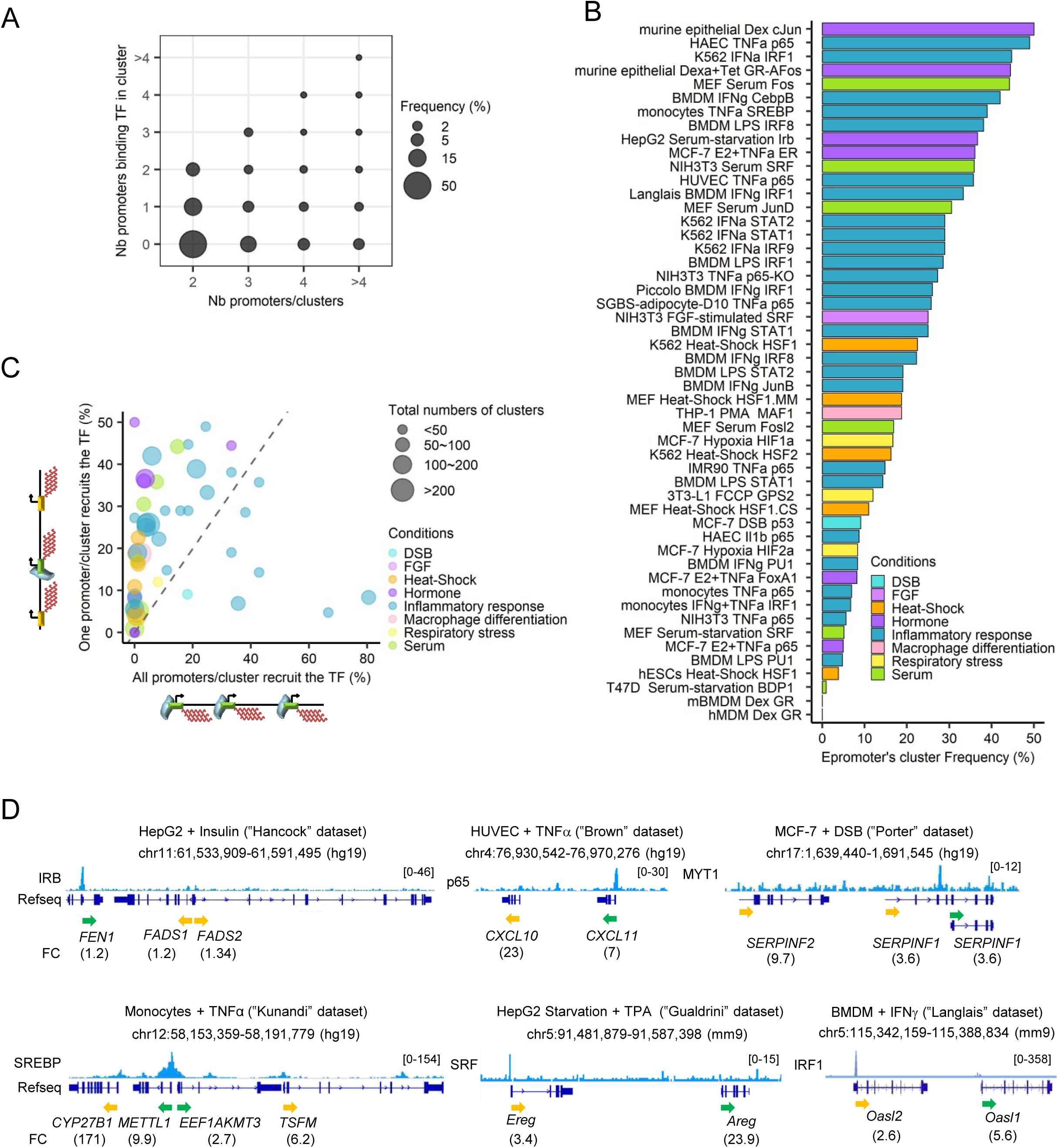
Frequency of Epromoter clusters in the different stress models. (A) Bubble plot showing the frequency of all the clusters identified in all the datasets by their number of promoters (x-axis) compared to their number of promoters recruiting their specific TFs (y-axis) determined by the pipeline. (B) Epromoter clusters frequency. Distribution of the Epromoter cluster frequency identified by the pipeline based on the distance for all the datasets. The colors indicated the different stimulation conditions of each dataset (light blue: double-strand breaks (DSB), light purple: Fibroblast growth factors (FGF) orange: heat shock, purple: hormone, blue: inflammatory response, pink: macrophage differentiation, yellow: respiratory stress, green: serum). (C) Distribution of the proportion of clusters where all clustered promoters recruit the key TF (x-axis) compared to the proportion of Epromoters-regulated clusters (y-axis). The size of the bubble corresponds to the number of total identified clusters by the pipeline. The colors indicated the different stimulation conditions of each dataset (light blue: double-strand breaks (DSB), light purple: Fibroblast growth factors (FGF) orange: heat shock, purple: hormone, blue: inflammatory response, pink: macrophage differentiation, yellow: respiratory stress, green: serum). (D) Examples of Epromoter-regulated clusters identified by the pipeline in different stress conditions. The stress dataset is indicated in top of each panel. The genomic tracks show the ChIP-seq signal of the TF used in each data after stimulation. The fold-change of induction is indicated below the name and orientation of the genes (Epromoter: green, co-induced genes: yellow).

### Systematic identification of stress-response clusters regulated by Epromoters

We applied our pipeline to systematically identify stress-response clusters regulated by Epromoters, using the 51 datasets described above (Table S2; Table S3). We identified a total of 606 non-redundant clusters potentially regulated by Epromoters. We distributed those clusters by the number of promoters per cluster in function of the number of promoters recruiting the associated TF (Fig. 3A). We found that amongst clusters harboring TF-bound promoters, 66% had only one promoter bound by the TF and 74% had fewer promoter bounds by the TF than the total number of promoters within the cluster (Fig. 3A). On average, we found 17 Epromoter-regulated clusters per dataset (Table S2). Out of 51 datasets, 49 contain at least one Epromoter cluster, ranging from 1 (“Ferrari” dataset (*49*), with serum starvation and the BDP1 TF) to 141 (“Phanstiel” dataset (*54*), with PMA and the MAF1 TF). To better assess the contribution of Epromoters to the regulation of stress-response clusters, we calculated the frequency of Epromoter-regulated clusters compared to all the clusters of induced genes found in each dataset (Fig. 3B; Supplemental Fig. S2). We found that Epromoter-predicted clusters represent between 0% and 50% of the identified clusters in each dataset, centered around 19% across all datasets. We then compared the proportion of Epromoter clusters to clusters where all promoters recruit the identified key TFs to the stress (Fig. 3C). We observed that very few clusters contained all their promoters recruiting the specific TFs, with a distribution centered at 4% of all clusters compared to 19% for Epromoter-like clusters. This points to a strong bias towards enrichment in stress-response clusters potentially regulated by Epromoters. Several examples of stress-response clusters potentially regulated by Epromoters are shown in Fig. 3D. Overall, our analyses suggested that regulation by Epromoters is a general mechanism in stress response rather than a specific or isolated phenomenon.

### Epromoters regulate gene clusters in different stress response models

We aimed to assess the role of Epromoters predicted by our pipeline on the expression of distal genes from the same cluster in their endogenous locus using CRISPR/Cas9 genomic deletion. We started by studying the *NUCB1* Epromoter cluster, which consists of six genes induced by HS in K562 cells (“Vihervaara” dataset) (Fig. 4A). The *NUCB1* Epromoter is the only promoter of the cluster recruiting HSF1 and HSF2 TFs (Fig. 4A). Consistently, the *NUCB1* Epromoter displayed enhancer activity in luciferase assays, while the *DHDH* promoter, which is the highest induced gene of the cluster, did not (Fig. 2G). Moreover, the *NUCB1* Epromoter significantly induced *DHDH* promoter activity after HS treatment of K562 cells as assessed by luciferase assay (Fig. 4B). Next, we tested the Epromoter regulation in a cellular context by using CRISPR-Cas9 mediated genomic deletion of the *NUCB1* Epromoter in K562 cells (Fig. 4C). The knockout of the *NUCB1* Epromoter resulted in the loss of *NUCB1* expression and a lack of induction of neighboring genes *TULP2* and *DHDH* in four homozygous ΔNUCB1 clones (Fig. 4D), demonstrating the enhancer activity of this regulatory element in the endogenous context. Note that the expression of the other three genes of the cluster (*PPP15RA*, *PLEKHA4,* and *GYS1*) were not induced in our conditions although their basal expression was slightly decreased in the knock-out cells (Fig. S3). Therefore, the promoter sequence of *NUCB1* is crucial for the regulation of the HS-induced gene *DHDH*.

**Fig. 4.**
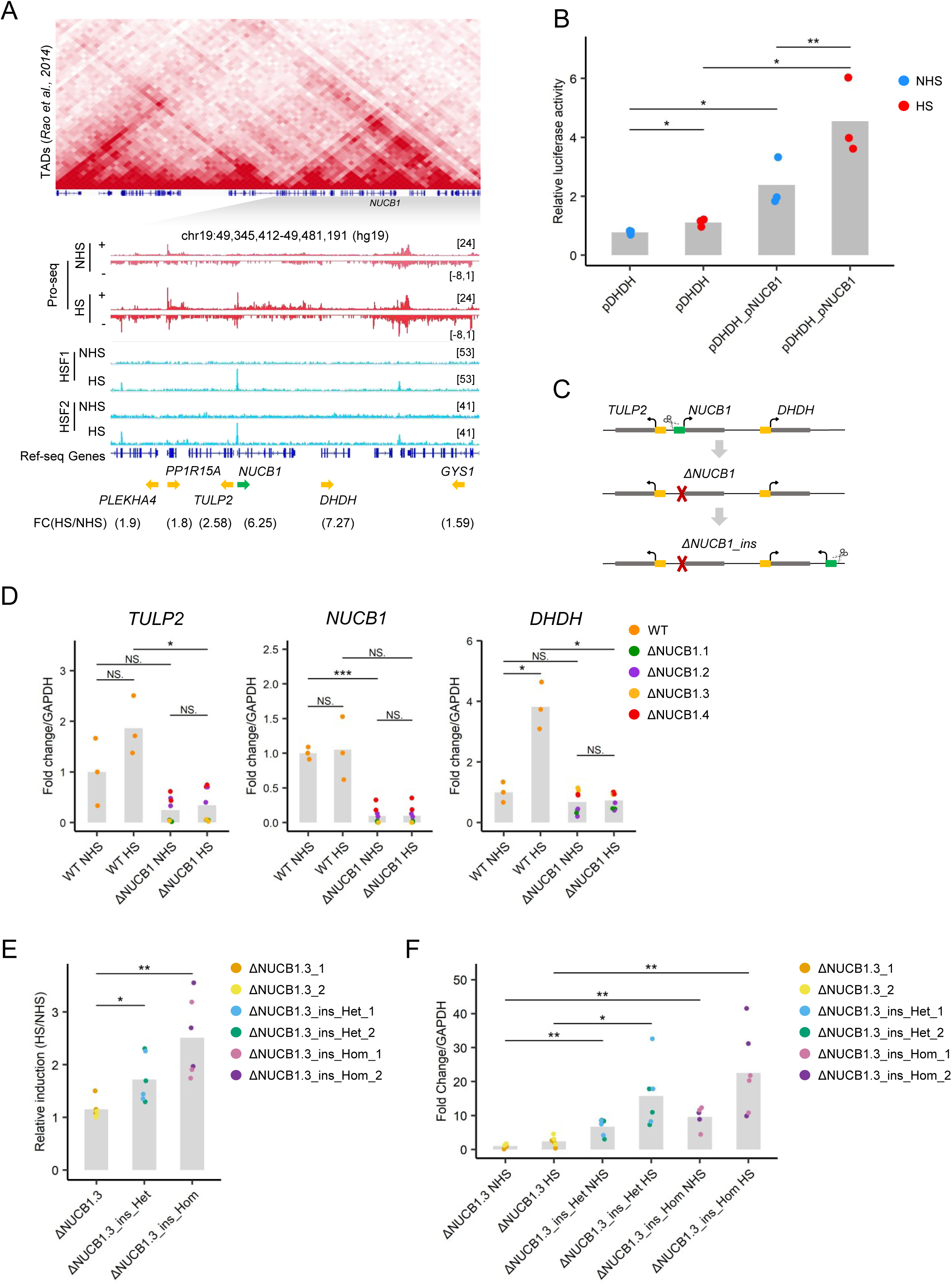
Epromoter-regulated cluster in the HS stress response. (A) The NUCB1 Epromoter-regulated cluster. (top) Hi-C triangular matrix (resolution 5kb) of the locus from K562 cells. The genomic tracks show the PRO-seq signal (in red) and HSF1 and HSF2 ChIP-seq signal (in blue) before or after HS from the “Vihervaara” dataset. The fold-change of induction is indicated below the name and orientation of the genes (Epromoter: green, co-induced genes: yellow). (B) Luciferase assays to quantify the promoter activity of the induced *DHDH* promoter with or without the *NUCB1* Epromoter acting as an enhancer before and after HS in K562 cells. *P* values were calculated by a paired one-sided Student’s t-test, \*\*\**P* < 0.001, \*\**P* < 0.01, \**P* < 0.1. (C) Schematic of the CRISPR-Cas9 genome deletion and re-insertion of the promoter region of the *NUCB1* Epromoter in K562 cells. (d) qPCR analysis of *TULP2*, *NUCB1*, and *DHDH* expression in wild type and 4 ΔNUCB1 mutants in K562 cells before and after HS. Values represent the relative expression of the samples normalized to the housekeeping gene *GAPDH* and compared to the unstressed wild-type cells. *P* values were calculated by a two-sided Student’s t-test, \*\*\**P* < 0.001, \*\**P* < 0.01, \**P* < 0.1. (E) qPCR analysis of the *DHDH* expression in 2 ΔNUCB1 mutants, 2 heterozygote ΔNUCB1-inserted mutants (green, blue), and 2 homozygote ΔNUCB1-inserted mutants in K562 cells before and after HS. Values represent the relative expression of the samples normalized to the housekeeping gene *GAPDH*, then compared before and after HS. *P* values were calculated by a two-sided Student’s t-test, \*\*\**P* < 0.001, \*\**P* < 0.01, \**P* < 0.1. (F) qPCR analysis of the promoter activity in 2 ΔNUCB1 mutants, 2 heterozygote ΔNUCB1-inserted mutants, and 2 homozygote ΔNUCB1-inserted mutants in K562 cells before and after HS. Values represent the relative expression of the samples normalized to the housekeeping gene *GAPDH* and compared to the mean of the ΔNUCB1 clones. *P* values were calculated by a two-sided Student’s t-test, \*\*\**P* < 0.001, \*\**P* < 0.01, \**P* < 0.1.

To test whether the *NUCB1* Epromoter can work as a distal regulatory element independently of its original genomic location, we re-inserted the *NUCB1* Epromoter sequence downstream of the *DHDH* gene in one of the ΔNUCB1 clones (Fig. 4C). We obtained two homozygous and two heterozygous cell lines with the re-inserted Epromoter. Upon comparing the induction of *DHDH* after HS between these modified cells and non-inserted cells (Fig. 4E), we observed that *DHDH* induction was restored in cells with the re-inserted *NUCB1*-Epromoter. We also measured the transcription downstream of the inserted sequence and observed HS-dependent transcriptional activity specifically in *NUCB1* re-inserted cells (Fig. 4F). This demonstrates that the *NUCB1* Epromoter sequence possesses intrinsic and location-independent enhancer and promoter activities.

In order to generalize our finding, we assessed the functional role of two additional predicted Epromoters induced by either TNFα or IFNγ cytokines. On the one hand, we used the “Lo” dataset (*51*), which includes RNA-seq and NFκB (p65) ChIP-seq data, both before and after 24 hours of TNFα stimulation in mouse NIH3T3 cells (Fig. 5A). Our pipeline identified 149 induced clusters, of which 8 were classified as Epromoter clusters, while none of the clusters harbor binding of p65 to all of the cluster-promoters (Fig. 5B, Table S2). Among these, we selected the *Cxcl1* Epromoter cluster for further investigation. This cluster comprises the potential Epromoter *Cxcl1* and its potentially regulated gene, *Cxcl2* (Fig. 5C). Both genes code for proteins of the CXC chemokine family and are strongly induced during the TNFα response. However, only the *Cxcl1* Epromoter is bound by the p65 TF. To explore the functional activity of this predicted Epromoter, we employed CRISPR-Cas9 genome editing to delete the *Cxcl1* Epromoter, generating three knock-out clones in NIH3T3 cells. The gene expression of *Cxcl1* and *Cxcl2* was then assessed following 4 hours of TNFα stimulation (Fig. 5D). The deletion of the *Cxcl1* promoter resulted in the complete loss of the *Cxcl1* expression but also in a significant reduction of the *Cxcl2* induction upon TNFα stimulation. These results indicate that the *Cxcl1* promoter exhibits enhancer activity on its neighboring gene *Cxcl2* during the TNFα response.

**Fig. 5.**
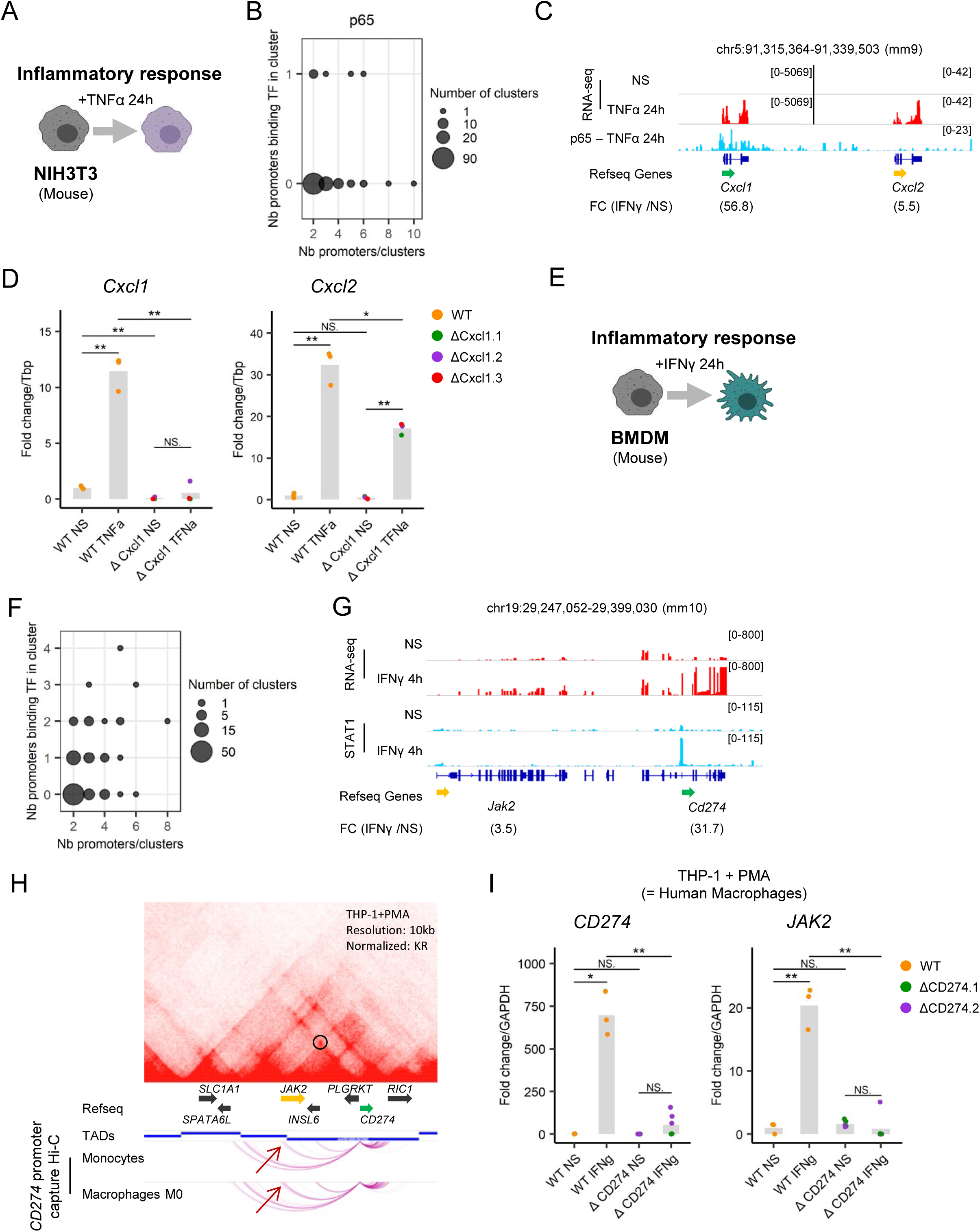
Epromoter-regulated cluster in the TNFα and the IFNγ stress response. (A) Schematic of the “Lo” dataset to induce TNFα response. (B) Bubble plot showing the number of all the clusters identified by their number of promoters (x-axis) compared to their number of promoters recruiting p65 (y-axis) determined by the pipeline. (C) Example of the *Cxcl1* Epromoter-regulated clusters identified by the pipeline in the TNFα stress response. The genomic tracks show the RNA-seq signal (in red) before or after 24 hours of TNFα stimulation and the p65 ChIP-seq signal (in blue) after 24 hours of TNFα stimulation from the “Lo” dataset. The fold-change of induction is indicated below the name and orientation of the genes (Epromoter: green, co-induced genes: yellow). (D) qPCR analysis of *Cxcl1*, *Cxcl2* expression in wild type (orange) and 3 ΔCxcl1 mutants (green, purple, and red) in the mouse NIH3T3 cells before and after stimulation TNFα for 4 hours. Values represent the relative expression of the samples normalized to the housekeeping gene *Tbp* and compared to the unstressed wild type. P values were calculated by a two-sided Student’s t-test, \*\*\**P* < 0.001, \*\**P* < 0.01, \**P* < 0.1. (E) Schematic of the experimental protocol from the “Piccolo” dataset to induce IFNγ response. (F) Bubble plot showing the number of all the clusters identified by their number of promoters (x-axis) compared to their number of promoters recruiting STAT1 (y-axis) determined by the pipeline. (G) Example of the *Cd274* Epromoter-regulated clusters identified by the pipeline in the IFNγ stress response. The genomic tracks show the RNA-seq signal (in red) and STAT1 ChIP-seq signal (in blue) before or after 4 hours of TNFα stimulation from “Piccolo” dataset. The fold-change of induction is indicated below the name and orientation of the genes (Epromoter: green, co-induced genes: yellow). (H) (top panel) Hi-C triangular matrix track (resolution at 10kb, Knight-Ruiz (KR) normalized) and the TAD from THP-1 cells. (bottom panel) Genomic interaction detected by CHiCAGO at the *Cd274* promoter by Promoter Capture Hi-C in monocytes, in vitro differentiated macrophages (M0). (I) qPCR analysis of *PDL1*, *JAK2* expression in wild type (orange) and 2 ΔPDL1 mutants (green and purple) in the human macrophages THP-1 cells with PMA before and after stimulation by IFNγ for 4 hours. Values represent the relative expression of the samples normalized to the housekeeping gene *GAPDH* and compared to the unstressed wild type. *P* values were calculated by a two-sided Student’s t-test, \*\*\**P* < 0.001, \*\**P* < 0.01, \**P* < 0.1.

On the other hand, we took advantage of the pipeline to test the model based on data from primary cells. We chose the “Piccolo” dataset (*34*), which consisted of RNA-seq and STAT1 ChIP-seq data before and after 4 hours of IFNγ stimulation in mouse Bone Marrow-Derived Macrophages (BMDM) primary cells (Fig. 5E). For instance, out of 103 distinct clusters, we identified 25 clusters potentially regulated by Epromoters (Fig. 5F, Table S2). From the identified clusters, we chose to focus on the *Cd274*-Epromoter cluster. This cluster includes the *Cd274* gene, coding for the ligand of the PD1 inhibitory receptor (PD-L1), and the *Jak2* gene, coding for the tyrosine kinase JAK2. Strikingly, both genes play an essential role in the inflammatory response and are often co-regulated (*59, 60*), however, only the *CD274* Epromoter binds the STAT1 TF (Fig. 5G). To experimentally validate the role of the *CD274*-Epromoter in the IFNγ response, we used PMA-induced in-vitro differentiated macrophage from the human THP-1 monocyte cell line, a classical model to study macrophage function. As expected, we observed that both *CD274* and *JAK2* genes were induced after 4 hours of IFNγ stimulation of *in vitro* differentiated THP-1 macrophages. Hi-C data from macrophage-differentiated THP-1 cells (*54*) revealed that both genes are in the same TAD. Consistently, analysis of Promoter Capture Hi-C interactions (*61*) centered on the *CD274* promoter showed an interaction with the *JAK2* promoter which increased from monocytes to macrophage differentiation (Fig. 5H). Deletion of the *CD274* promoter in THP-1 cells resulted in significant impairment of both *CD274* and *JAK2* induction after 4 hours of IFNγ stimulation in *in vitro* differentiated THP-1 macrophages (Fig. 5I). These findings indicate that the *CD274* promoter functions as an enhancer for *JAK2* in THP-1 differentiated cells. Overall, these results demonstrated the robustness of the pipeline in predicting *bona fide* Epromoters across various stress responses, and predictions from primary cells.

### Disruption of Epromoter-bound SRF TF impact on the expression of the whole clusters

We observed that for Epromoter-regulated clusters, only one of the promoters is bound by the key TF that is required for the stress response. This implies that disruption of this TF should impact the expression of all the genes in the cluster, therefore reflecting the regulatory role of the Epromoter. To address this question, we used the “Esnault” dataset (*31*) of NIH3T3 mouse fibroblasts upon 30 min of serum stimulation after starvation (Fig. 6A). Serum response induced a rapid gene induction, which is mainly dependent on the binding of the Serum Response Factor (SRF) (*62*). The pipeline identified 53 clusters, of which 19 were classified as Epromoter clusters (Fig. 6B; Table S2). Additionally, the study included RNA-seq data in conditions of SRF inhibition. Examination of the *Msrb3* Epromoter-like cluster (Fig. 6C) revealed that SRF inhibition abolished the induction of the *Msrb3* gene and the *Lemd3* co-induced gene located in the same cluster (Fig. 6D), suggesting that SRF binding on the *Msrb3* Epromoter is also required for the induction of the *Lemd3* gene. To generalize this observation, we quantified the effect of SRF inhibition on the serum induction of all Epromoter-genes and co-induced genes. As shown in Fig. 6E, the induction of both sets of genes was similarly affected by SRF inhibition. This finding supports a dependence of co-induced genes on Epromoters for TF recruitment and regulatory function.

**Fig. 6.**
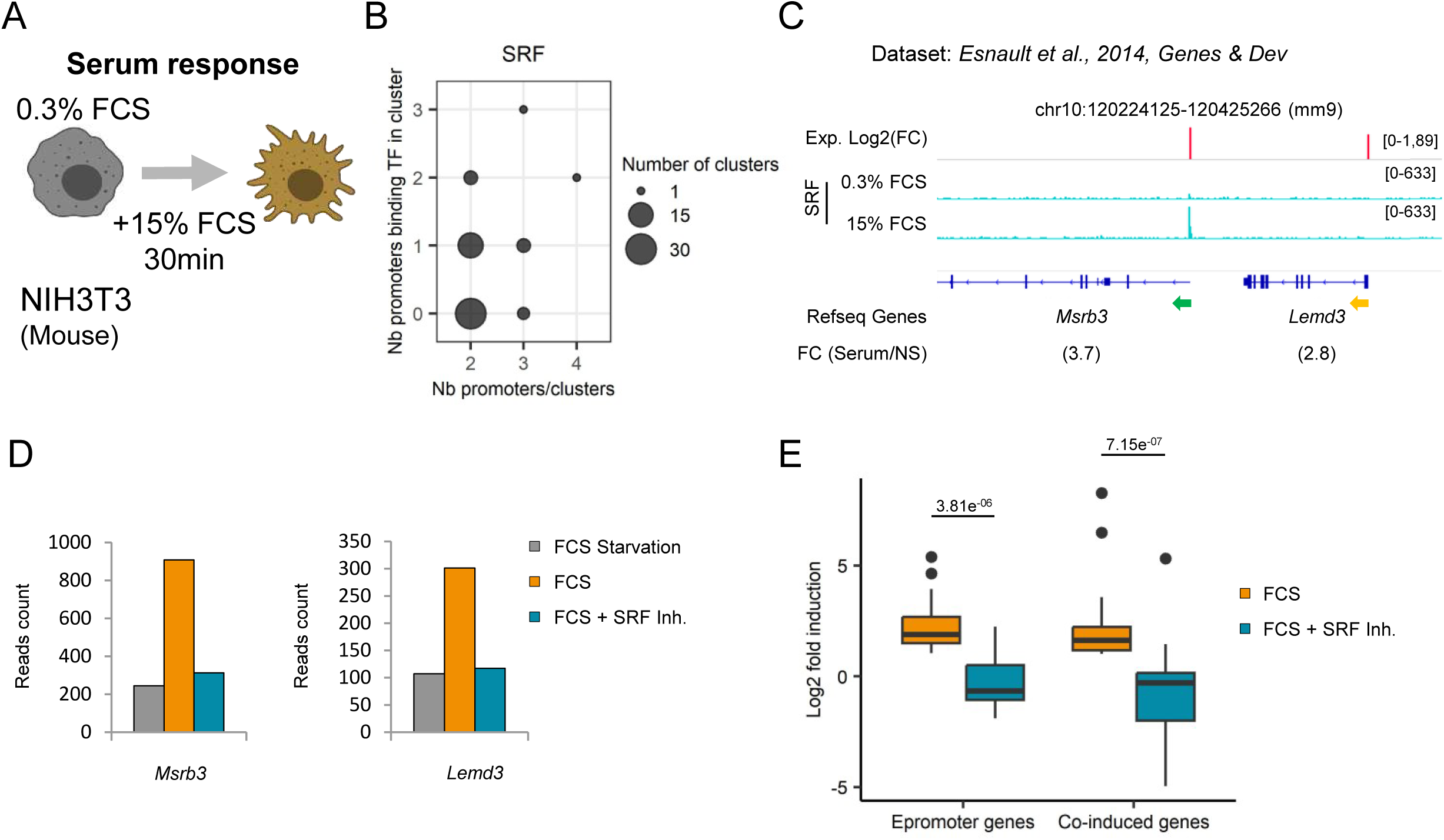
TF disruption at Epromoter-regulated clusters in the serum response. (A) Schematic of the “Esnault” dataset to induce serum response (15% FCS) after serum starvation (0.3% FCS). (B) Bubble plot showing the number of all the clusters identified by their number of promoters (x-axis) compared to their number of promoters recruiting SRF (y-axis) determined by the pipeline. (C) Example of the *Msrb3* Epromoter-regulated clusters identified by the pipeline in the serum stress response. The genomic tracks show the log2 fold-change of the RNA-seq signal (in red) and SRF ChIP-seq signal (in blue) before or after serum stimulation from the “Esnault” dataset. The fold-change of induction is indicated below the name and orientation of the genes (Epromoter: green, co-induced genes: yellow). (D) RNA-seq reads count of the genes *Msrb3* and *Lemd3* after starvation (grey, FCS Starvation), stimulated by serum (orange, FCS) or stimulated by serum with inhibited SRF (blue, FCS+ SRF inh). (E) Boxplot of the fold of induction (in log2) of the Epromoters and co-induced genes identified by the pipeline after FCS stimulation (orange) or stimulation with inhibited SRF (blue). *P* value was calculated by a paired two-sided Wilcoxon’s test.

## Discussion

To better understand the regulation of stress-response genes, we analyzed a series of published datasets from different proteotoxic, environmental and inflammatory stresses, including gene expression data and ChIP-seq for key transcription factors involved in the same stress responses. We initially found that stress-response genes are frequently located in close genome proximity, pointing to shared regulatory networks. We then aimed to assess the implications of Epromoters in the regulation of stress-response clusters. We previously showed that IFNα-induced Epromoters preferentially recruit the key interferon response complex (STAT1/STAT2, IRF1/IRF9) and regulate the interferon response of neighboring genes (*18*). Leveraging these findings, we developed a pipeline whereby stress-response clusters potentially regulated by Epromoters were identified based on the fact that only one of the induced promoters (i.e., the Epromoter) recruited the key TF involved in the regulation of the stress response. We observed frequent stress-response clusters potentially regulated by Epromoters in the majority of the studied stress models. We systematically validated the predicted enhancer activity of HS-induced Epromoters and confirmed the regulatory role of Epromoters on stress-induced clusters within their endogenous loci, across three different stress conditions and cell types. Furthermore, using the HS-inducible *NUCB1-DHDH* loci we demonstrated that relocating an Epromoter within its genomic locus did not affect its function, proving that the Epromoter sequence is sufficient to function independently of its original genomic position. Overall, our findings indicate that Epromoters-regulated clusters are a general mechanism of stress response where Epromoters specifically recruit key stress-associated TFs, thereby facilitating the rapid and coordinated activation of genes within such clusters to have a more efficient response to intra- and extracellular stress signaling.

One limitation of our study regards the potential underestimation of stress-response clusters regulated by Epromoters. First, our current pipeline identified clusters with a threshold at 100 kb between the TSS of co-induced genes. More optimal approaches might use 3D interaction data or an adaptive threshold distance specific for each stress condition. Second, we considered Epromoter-regulated clusters as only those in which only one promoter recruits the key TF. However, more complex situations also exist where only a limited number of promoters do not recruit the TF (e.g., *GBP* cluster; Fig. 4A). For instance, we previously identified the IFIT cluster composed of a family of 4 IFNα-induced genes with 3 recruiting the STAT1/2 and IRF1/9 factors (*18*). The deletion of one of the promoters (*IFIT3*) inhibited the induction of the fourth gene (*IFIT2*). Finally, our identification of Epromoter clusters was also limited by the availability of stress-related datasets. The automated pipeline we have developed will ease the systematic identification of new Epromoters clusters in the future.

The impact on neighbor gene expression after deletion of the Epromoter could, in theory, be explained by alternative mechanisms, including regulatory function of the gene product and readthrough effects. In previous studies we have shown that re-expression of Epromoter-associated genes did not rescue the expression of neighbor genes (*15, 18*), indicating direct regulation of neighboring gene expression. Various types of stress, such as high temperatures and viral infections, can induce Pol II transcription past the normal termination site into neighboring genes, a process termed readthrough transcription (*2*). However, this mechanism will only work when the neighbor gene is located downstream the stress-response gene in a tail-to-head configuration. Anyhow, the rescue of *DHDH* induction observed after repositioning of the HS-Epromoter *NUCB1* are not compatible with the two aforementioned mechanisms and rather support a direct regulatory role of Epromoters.

Stress response is a cellular mechanism that allows cells to adapt to various environmental disturbances that disrupt their homeostasis. Numerous factors can be considered as stress factors. The most well-known stressors include DNA-damaging agents, nutrient deprivation, temperature fluctuations, chemical toxins, and infectious agents. Each type of stress triggers a specific response that includes rapid changes in gene transcription. Even though the transcription landscape can be greatly modified, such changes are mostly mediated by a few and very specific TFs. The implications of a small number of TFs upon stress allow the cells to quickly respond and adapt to it but also to swiftly return to normal function once the stress is alleviated. Previous studies suggested that Epromoters might be involved in stress response and, in particular, the interferon response (*15, 18, 20, 25*). For instance, active Epromoters in HeLa cells, which have a constitutive type 1 interferon response, are significantly associated with interferon response (*15*). Indeed, inhibition of type I interferon in HeLa cells preferentially affects the activity of Epromoters as compared with distal enhancers (*18, 20, 25*). Moreover, using high-throughput reporter assays, we previously showed that IFNα-induced genes are frequently organized in clusters regulated by a single Epromoter (*18*). Within these clusters, Epromoters function as regulatory hubs that selectively recruit TFs essential for IFNα-driven responses, coordinating gene expression across the cluster. Our current study validates and extends these findings by demonstrating a predominant role of Epromoters in the regulation of stress-response genes in general.

Cell identity genes are known to be regulated by distal enhancers, often organized in clusters of enhancers in large intergenic regions (*63*). In contrast, housekeeping genes lack interactions with distal enhancers but are regulated by promoter-promoter interactions mediated by the binding of the Ronin TF (*26*). Here, we propose that stress-response genes are preferentially organized in clusters in which often one of the promoters plays a primordial role in the coordination of all genes of the cluster. Thus, our findings demonstrate that clustering of gene regulatory elements, whether enhancers or promoters, is a general principle of gene regulation that is not limited to cell identity or housekeeping genes but extends to stress and inflammatory response genes that are essential for cellular homeostasis.

It is clear that not all stress-response genes are found in clusters. While unclustered genes are more likely to be regulated by typical enhancers, our results suggest that clustered genes are preferentially regulated by Epromoters. This is supported by our previous study showing that unclustered interferon-induced genes are associated with distal regulatory elements binding ISGF3 complex, while those found within clusters are more frequently associated with ISGF3-bound Epromoters (*18*). In line with these observations, visual inspection of the epigenetic and TF binding profiles at typical stress response clusters, suggests that no potential distal enhancers are found within the TADs containing the clusters.

Features defining the enhancer *versus* promoter activity of regulatory elements are a fundamental question in the gene regulation field and a focus of extensive research (*13, 19, 63–69*). In this context, what are the mechanistic bases leading to the enhancer activity of stress-response Epromoters? We previously showed that constitutive Epromoters bind a higher number of TFs and harbor a more complex combination of TF binding sites as compared with classical promoters (*15*), suggesting that one of the potential mechanisms mediating enhancer function might be the efficient recruitment of key TFs. Here, we observed that HS-induced Epromoters harbor a higher density of HSF1/2 motifs as compared with other induced promoters, which could explain the highly efficient recruitment of these TFs. Similarly, the density and quality of interferon-response elements are higher at interferon-induced Epromoters as compared to the promoters of other induced genes, but similar to the predicted distal enhancers (*18*). This is therefore likely that a high density of stress-response binding sites can lead to increased efficiency of TF recruitment and/or stabilization required for the enhancer function of Epromoters.

The regulation of stress-response clusters by Epromoter aligns well with the notion of inducible condensates defining discrete membrane-less sub-nuclear compartments containing a high concentration of RNA polymerase and key TFs and where efficient transcription can be triggered (*2, 7, 27*). A striking example is provided by NFkB-regulated genes in response to TNFα stimulation (*70*). Experimental removal of a gene from the NFkB-dependent multi-gene complex was shown to directly affect the transcription of its interacting genes, suggesting that co-association of co-regulated promoters might contribute to a hierarchy of gene expression control. Similarly, it is likely that Epromoters might tend to generate small, localized hubs due to their higher recruitment of TFs. In this context, the highly efficient recruitment of stress-response factors observed at Epromoters could lead to the formation of condensates through the interaction of TF’s intrinsic disordered regions, as seen with HSF1 (*71*) and RNA Pol II (*72*). This, in turn, could either facilitate the assembly or maintenance of stress condensates by tightening promoter-promoter interactions or bringing specific transcriptional regulators required for the regulation of the neighbor genes. In support of this hypothesis, we found that the *CD274* and *JAK2* coregulated genes are within the same interacting loop in unstimulated macrophages, but the interaction is increased upon IFNγ stimulation, in agreement with previous results suggesting that a pre-establish loop between *cis*-regulatory elements can facilitate rapid gene induction (*43, 73*). This would be particularly relevant in the case of rapid and coordinated regulation of gene expression in response to environmental or intrinsic cellular stimuli (*2*). Our work, thus, provides a framework for future studies aiming to address the contribution of Epromoters in the formation or stabilization of transcriptional condensates in response to different stress signaling.

Our findings have significant implications for the understanding of the evolution of regulatory elements and stress-response clusters. On the one hand, many of the identified stress-response clusters are likely to arise through gene duplication events. Indeed, many of the identified clusters consist of genes from the same gene family, including, for example, *OAS, IFIT*, *CXCL*, *CXCR* and *GBP* clusters (Table S3). For instance, gene expansion has been shown to significantly contribute to the evolution of the IFN system and suggested to confer a selective advantage to the host species (*74*). It is possible that during the duplication of stress-response genes, ancestral promoters’ elements have acquired enhancer functions to coordinate the response of the new gene isoforms within the cluster. On the other hand, recent works suggested that repurposing of promoters and enhancers facilitated regulatory innovation and the origination of new genes during evolution (*5, 67, 75–80*). Similarly, the proximity to an Epromoter-associated locus might provide a rapid co-option for neighbor genes to acquire new functions in stress response. Indeed, we observed that for HS clusters, Epromoter genes were directly associated with HS-related functions, while no significant enrichment was found for the distal Epromoter-regulated genes. These observations are reminiscent of previous findings that rapid induction of immediate-early genes in response to growth factor stimulation is accompanied by co-upregulation of their neighboring genes (*6*). These observations suggest that transcriptional activation has a ripple effect, which may be advantageous for coordinated expression. It is plausible that these genes, while initially unrelated to the stress responses, may have been co-opted into stress-related pathways by Epromoter hijacking. Alternatively, it is equally possible that some of the distally co-regulated genes might be induced by the proximity to the Epromoters without any physiological relevance. Future work will need to be performed to more systematically assess the functional role of the distal-associated genes.

One way to explore potential Epromoter hijacking contributing to functional co-option is by examining large-scale structural variations across species. For instance, Gilbertson et al. (*81*) described a chromosomal inversion of approximately 8 Mb on chromosome 5 between mice and humans. In mice, the genes *Oasl1* and *Oasl2* are separated by this 8 Mb region from the *P2rx7* gene, whereas in humans, the inversion brings the *OASL* gene (the ortholog of *Oasl1* and *Oasl2*) within just 100 kb of *P2RX7*. This genomic rearrangement appears to have regulatory consequences as both *OASL* and *P2RX7* are co-induced by LPS stimulation in humans, while in mice, *Oasl1* and *Oasl2* are induced by LPS, but not *P2rx7*. We identified *Oasl2* as an Epromoter for *Oasl1* in mouse macrophages in response to IFNγ (*50*) (Fig. 3D). Interestingly, we noted that *P2rx7* is not induced in mice by IFNγ, while in human macrophages, both *OASL* and *P2RX7* are induced by IFNγ (Fig. S4B; expression data from (*82*)). These observations likely reflect the co-optation of *P2RX7* into the inflammatory response by hijacking the *OASL* Epromoter.

As dysregulation of stress response signaling pathways is involved in multiple pathologies, it is likely that genetic or epigenetic alterations of stress-Epromoters might have a pleiotropic role in the etiology of inflammatory- and stress-related diseases (*2, 83*). Several studies, including ours, have demonstrated that human genetic variation within Epromoters influences distal gene expression (*15, 21–24*). Moreover, specific examples highlight the distal impact of disease-associated variants within Epromoters (*19, 84–90*). As Epromoters potentially control several genes at the same time and efficiently recruit key TFs, mutations in these regulatory elements are expected to have a stronger pathological impact, as compared to typical promoters. This might result from the regulation of multiple genes either involved in the same (additive or synergistic effects) or different (pleiotropy) pathways. Future work will require a systematic association of Epromoters with genetic traits coupled with functional studies of the target genes in order to demonstrate their involvement in the context of stress-related disease.

All in all, by leveraging extensive genomic and functional datasets, our study explores the relationship between Epromoter and the regulation of stress-response clusters, shedding light on the complex regulatory mechanisms leading to rapid gene induction during the cellular responses to intra- and extracellular stress signaling. Given the involvement of stress response in pathological conditions and the proposed pleiotropic role of Epromoters (*19*), we foresee an important contribution of this mechanism to disease.

## Methods

### Selection of Stress-related datasets

Selection of stress-related datasets was achieved by manual inspection of the literature. We selected datasets for which differential gene expression in any stress conditions from mouse or human and ChIP-seq for key TFs involved in the same stress conditions was available. All stress-related datasets used in this study are listed in the Table S1 and described in the Table S2.

### Distance distribution of induced genes

Induced genes were identified from gene expression data derived from stress response datasets (Table S2), considering only protein-coding genes. Reference gene annotation files were downloaded from the UCSC Genome Browser: human (hg19: https://hgdownload.soe.ucsc.edu/goldenPath/hg19/database/refGene.txt.gz, hg38: http://hgdownload.soe.ucsc.edu/goldenPath/hg38/database/refGene.txt.gz), and mouse (mm9: https://hgdownload.soe.ucsc.edu/goldenPath/mm9/database/refGene.txt.gz, mm10: https://hgdownload.soe.ucsc.edu/goldenPath/mm10/database/refGene.txt.gz). The distances between the transcription start sites (TSS) of induced genes were calculated using Bedtools with the following parameters: bedtools closest -a $file -b $file -d -io -t first. For comparison, random gene sets containing the same number of genes as the induced gene sets were generated by randomly selecting protein-coding genes from the annotation files. The distances between the TSS of these random genes were calculated in the same manner as for the induced genes. To assess statistical significance in the distance distributions between induced and random genes, a Kolmogorov-Smirnov (K-S) test was performed. To quantify the deviation between the distance distributions of induced and random genes, deviation scores were defined using a Q-Q plot-based method. Specifically, the Euclidean distances between the observed induced-random points (divided into 1000 quantiles) and the expected line of equality (y = x, representing no difference between distributions) were calculated. The sum of these Euclidean distances was defined as the deviation score.

### Identification of Epromoter-regulated clusters

We developed a pipeline to identify Epromoter-regulated clusters using expression data (RNA-seq, Pro-seq or microarrays) and ChIP-seq for key transcription factors involved in the same stress responses. The pipeline and its’ usage can be accessed from Github: https://github.com/Spicuglia-Lab/Epromoter-like-cluster-pipeline. The pipeline identifies clusters of induced genes in stimulated conditions on the basis of: (i) distance between two induced gene’s transcription start site (TSS) must be less than 100 kb, and (ii) genes per cluster should be more than 1. TF binding peaks overlapping with the TSS-proximal regions (±1kb) is considered as transcription factor is binding to the promoter of the gene. When there is only one promoter binding with TF in an induced gene clusters, this promoter is considered as Epromoter. For each execution of pipeline, it takes differentially expressed genes and TFs binding peaks as input. The input data requires three files: differentially expressed genes in stimulated condition with fold change, TF binding peaks in the same condition, reference transcript annotation file. The output of pipeline includes two files: (1) gene clusters with TF binding information and (2) Epromoter-regulated clusters. Additionally, the pipeline provided a TAD (Topologically associating domain) clustering method which consider induced genes working as clusters in a TAD. TAD clusters where only one promoter was bound by the TF were selected as Epromoter-regulated clusters. The pipeline was coded with Python 3.7.12 and implemented in Linux environment. The dependent libraries include pandas, plotnine, numpy, subprocess, seaborn, itertools, matplotlib, scipy, pybedtools. The list of Epromoter-regulated clusters for all datasets is provided in the Tables S3.

### Cell culture

Cell line K562 (CCL-243), a chronic myelogenous leukemia cell line, was obtained from the ATCC (American Type Culture Collection) and maintained in RPMI 1640 media (Thermo Fischer Scientific) supplemented with 10% FBS (Thermo Fischer Scientific) at 37°C and 5% CO2. Cells were passed every 3 days at 2 × 10^5^ cells/mL and routinely tested for mycoplasma contamination. Cell line NIH3T3 (CRL-1658), an embryonic mouse fibroblast, and maintained in DMEM media (Thermo Fisher Scientific) supplemented with 10% FBS (Thermo Fischer Scientific) at 37°C and 5% CO2. Cells were passaged every 2 days at 2 × 10^5^ cells/mL and routinely tested for mycoplasma contamination. THP-1, a human acute monocytic leukemia cell line, was obtained from DSMZ (ACC 16). Cells were grown in RPMI 1640 media (Thermo Fischer Scientific) supplemented with 10% FBS (Thermo Fischer Scientific) at 37°C and 5% CO2. Cells were passaged every 3 days at 106 cells/mL and routinely tested for mycoplasma contamination.

### Stress conditions

#### Heat-shock treatment

Exponentially growing K562 cells were placed in water baths at 37°C (NHS) or at 43°C (HS) for 1 hour followed by a recovery step of 2 hours in an incubator at 37°C and 5% CO2.

#### TNFα treatment

Exponentially growing NIH3T3 cells were stimulated by adding one volume of fresh media containing 10 ng/mL TNFα (Abcam, ab9642), for a final concentration of 5ng/mL, during 4 hours.

#### IFNγ treatment

To induce macrophage differentiation, THP-1 cells were firstly plated on 6-well plates (2 × 106 cells/well), in media containing 10 ng/ mL phorbol 12-myristate 13-acetate (PMA; Sigma-Aldrich, P1585) for 48 h. After 48 h of incubation, the PMA-containing media was replaced with fresh media, and cells were incubated for an additional 24 h. THP-1 *in vitro* differentiated macrophages were then stimulated by replacing media with 2 mL of fresh media containing 100 ng/mL of IFNγ (Miltenyi Biotec, 130-096-484) for 4 hours, upon which the extraction of RNA was performed. For each group, three independent simulations were made.

### Luciferase reporter assay

The pGL4-SV40 vector was created by cloning the promoter SV40 into the pGL4.12 vector (Promega, #E6671) at the BglII and HindIII restriction sites. The human promoters (500 bp) tested were cloned into the pGL4-SV40 vector downstream the luciferase gene at the SalI and BamHI restriction sites (Table S4). The human promoters *HSPA1A* and *DHDH* were cloned into the pGL4.12 vector (Promega, #E6671) at the BglII and HindIII restriction sites. The human promoter *DHDH* was also cloned at the BglII and HindIII restriction sites instead of the SV40 promoter in the pGL4-SV40-NUCB1 vector (Table S4). For cell transfection, 1 × 10^6^ K562 cells were mixed with 1 μg of each construct and 125 ng of renilla using the Neon™ Transfection System (Thermo Fisher Scientific; 100µl tips; pulse voltage 1450 V, pulse width 10, pulse number 3) and cultured in 1.2 mL in 12-well plates. For heat shock treatment, 24 h after transfection cells were placed in water baths at 37°C (NHS) or at 42°C (HS) for 1 hour followed by a recovery step of 2 hours in an incubator at 37°C and 5% CO2. The Dual-Luciferase kit (Promega, E1980) was used to measure luciferase and renilla luminescence following the manufacturer’s recommendations. Data were normalized to renilla values and represented as the fold-change of relative light units over the pSV40-luc vector. Experiments were performed in triplicate.

### Motifs enrichment analysis

The HSF1 and HSF2 binding sites in the human genome were recovered from the JASPAR database (MA0486.2 for HSF1 and MA0770.1 for HSF2) (Sandelin, 2004) and overlapped with the promoter of all the induced genes induced by the HS response according to the PRO-seq data from the “Vihervaara” dataset or random promoters. The promoter coordinates were extended from −1250bp to +750bp centered on the TSS. The overlap was accomplished by using the function intersect of bedtools with the overlaps count (-c) between the coordinates of the HSF1/HSF2 JASPAR motifs and the coordinates of all the induced genes.

### Functional enrichment

GO enrichment in biological processes was assessed using g:Profiler (*91*) and default options. For the statistical background, we used the list of all genes associated with the induced promoters. Enrichment scores were calculated using the g:GOSt native method. The R package “rrvgo” was used with default options to concatenate the GO biological process term (*92*).

### CRISPR/Cas9 genome deletion

For promoter deletion we used the web tool CrispRGold and the CRISPR-Cas9 guide RNA from IDT to design two guide RNAs flanking the Epromoter regions. For each Epromoter, two primers were designed flanking the target region that allows us to identify the wild-type and the mutant alleles. We used the Alt-R CRISPR-Cas9 System from IDT (Integrated DNA Technology) where Ribonucleoprotein (RNP) complexes were assembled in vitro. The transfection was made with 5 × 10^5^ cells, 1 µl of each RNP complex, and 2 µl of Alt-R® Cas9 Electroporation Enhancer (IDT, 075915) using the Neon™ Transfection System (Thermo Fisher Scientific; 10µl tips; pulse voltage 1450 V, pulse width 10, pulse number 3) and cultured in 1.2 mL into the K562 and THP-1 cells. For the deletion of the Epromoter *Cxcl1* in NIH3T3 cells, we used CRISPRIMAX lipofectamine (Invitrogen, CMAX00001) following a protocol from IDT (Alt-R CRISPR-Cas9 System: Cationic lipid delivery of CRISPR ribonucleoprotein complexes into mammalian cells). Three days after transfection, cells were cultured in 5 × 96-well plates at limited dilution (0.3 cells/100 μL/well). After 2–4 weeks the clones were screened for homologous deletion using the kit Phire Tissue Direct PCR Master Mix (Thermo Fisher Scientific, F170L). Clones with homologous deletion were those showing a mutant band of the expected size and no wild-type band. The clones were then validated by Sanger sequencing. Primers and gRNAs are listed in Table S4.

### CRISPR/Cas9 genome insertion

For the insertion of the *NUCB1* Epromoter in K562, we used the web tool CRISPR-Cas9 guide RNA from IDT to design one guide RNA. Four primers were designed flanking and inside the inserted region that allows us to identify the wild-type and the mutant alleles. We created the knock-in insert by combining two homology arms flanking the *NUCB1* Epromoter and then used the Guide-it Long ssDNA Production System v2 (Takara, 632666) to produce a long single-stranded DNA. Transfection was performed using the Alt-R CRISPR-Cas9 System (IDT). The transfection was made with 5 × 10^5^ cells, 1µL of RNP complex, 1µg of ssDNA insert, 2µl of Alt-R® Cas9 Electroporation Enhancer (IDT, 075915), and 5µl of Alt-R™ HDR Enhancer V2 (IDT, 10007910) using the Neon™ Transfection System (Thermo Fisher Scientific; 10µl tips; pulse voltage 1450 V, pulse width 10, pulse number 3) and cultured in 1.2 mL into the Δ3*NUCB1* K562. Three days after transfection, cells were cultured in 5 × 96-well plates at limited dilution (0.3 cells/100 μL/well). After 2–4 weeks the clones were screened for homologous deletion using the kit Phire Tissue Direct PCR Master Mix (Thermo Fisher Scientific, F170L). Clones with homologous insertion were those showing a mutant band of the expected size and no wild-type band. Clones with heterozygous insertion were those showing a mutant band and a wild-type band of the expected size. The clones were then validated by targeted Long-read sequencing. Primers and gRNAs are listed in Table S4.

### Gene expression analysis

RNA was extracted using the RNeasy Plus Mini Kit (Qiagen, 74106) according to the manufacturer’s instructions with DNase treatment. 1 to 2.5 µg of total RNA was reverse transcribed using the SuperScript™ VILO™ (Thermo Fisher Scientific, 1755250). qPCR reactions were made using the SYBR green Master Mix (Thermo Fisher Scientific, 4367660) and the measurement was made using the Applied Biosystems QuantStudio 6 Flex Real-Time PCR System. 25ng of cDNA was used for the qPCR. The relative expression was analyzed using the relative standard curve method. The *GAPDH* gene was used for normalizing samples in humans and the *Tbp* gene was used for normalizing samples in mice. Each point on the figures corresponds to an independent RNA and cDNA preparation. The mean values were calculated between the relative expression values of the biological replicates. The values were then normalized either by the unstimulated WT or by the unstimulated cells depending on the experiment. To assess genomic DNA degradation, genomic primers targeting the *PAX5* promoter sequence were used. Primers used are listed in Table S4.

### 3D chromatin organization of the *CD274* locus

Hi-C data in THP-1+PMA cells were retrieved from (*54*). TADs called from Hi-C experiments (HindIII) THP-1 cells were taken from (*93*). DNA interactions centered on the *CD274* promoter with the associated CHiCAGO score were taken from published Promoter Capture Hi-C (*61*). All 3D chromatin interaction data were visualized using the New WashU Epigenome Browser (epigenomegateway.wustl.edu) (*94*).

### TF disruption analysis

RNA-seq data before and after serum stimulation (0.3% FCS and 15% FCS respectively) and with SRF inhibition were obtained from (*31*). SRF inhibition was achieved using the specific pathway inhibitors Latrunculin B (LatB) and U0126. From the serum-induced genes, the pipeline identified the Epromoters-regulated clusters from which the genes were classified depending on their promoters (Epromoter or co-induced). For each gene, the induction fold change was calculated by comparing read counts from serum-stimulated (15% FCS) and serum-starved (0.3% FCS) conditions, both with and without SRF inhibition.

### Statistical analysis

All statistical tests were performed using RStudio (2022.02.0+443), under version of R (4.1.3) and are indicated in the legend of each figure.

## Data availability

All publicly available datasets are listed in the Supplemental Table S1. Processed data for all datasets is provided in Zenodo (doi:10.5281/zenodo.14211588).

## Code availability

Custom scripts used in this study are available at GitHub (https://github.com/Spicuglia-Lab/Epromoter-like-cluster-pipeline).

## Acknowledgments

Work in the laboratory of Salvatore Spicuglia was supported by the ANR (grant numbers ANR-23-CE12-0008-01 AND ANR-20-CE12-0023). JM was supported by a PhD fellowship from the French ministry of research and a *LIGUE contre le Cancer* fellowhip. JW was supported by PhD fellowships from the EU-funded Innovative Training Network “Molecular Basis of Human Enhanceropathies” (Horizon 2020 research and innovation program under Marie Sklodowska-Curie grant agreement no. 860002) and from the Marseille Institute of Rare Diseases (MARMARA). GF was supported by a PhD fellowship from the *LIGUE contre le Cancer*.

## Contributions

SS and JM designed the study. HS and JW developed the bioinformatic pipeline. JM conducted most experimental work and performed bioinformatics analyses. CS and GF performed experimental validations. SS and JM analyzed the results and wrote the manuscript. All authors read and revised the manuscript.

## Supplementary legends

**Fig. S1.**

The top panels display the distance distribution of the induced genes (colors) compared to the same numbers of random genes (grey) by dataset. The bottom panels display the deviation score calculated between induced gene distance and random gene distance for each dataset. *P* values were calculated by Kolmogorov–Smirnov (KS) test.

**Fig. S2.**

(A) Example of the *HSPA4* Epromoter-regulated cluster identified specifically by the pipeline using the TAD clustering method in the HS stress response. The genomic tracks show the PRO-seq signal (in red) and HSF1 and HSF2 ChIP-seq signal (in blue) before or after HS from the “Vihervaara” dataset. The topological associated domains (TAD) from K562 are on the top. The fold-change of induction is indicated below the name and orientation of the genes (Epromoter: green, co-induced genes: yellow).

(B) Example of the *HSPB1* Epromoter-regulated cluster identified specifically by the pipeline using the distance clustering method in the HS stress response. The genomic tracks show the PRO-seq signal (in red) and HSF1 and HSF2 ChIP-seq signal (in blue) before or after HS from the “Vihervaara” dataset. The topological associated domains (TAD) from K562 are on the top. The fold-change of induction is indicated below the name and orientation of the genes (Epromoter: green, co-induced genes: yellow).

(C) Luciferase assay to quantify the promoter activity of the control *HSPA1A* promoter before (blue) and after (red) HS in the K562 cells. The results were normalized on the renilla activity. *P* value was calculated by a paired one-sided Student’s t-test, \*\*\**P* < 0.001, \*\**P* < 0.01, \**P* < 0.1.

(D) Scatterplot of the luciferase signal to quantify the enhancer activity of predicted Epromoter before (x-axis) and after (y-axis) HS. The predicted Epromoter (green) and the induced promoters (yellow) are highlighted

(E) Significantly enriched GO biological processes for the HS-induced genes in K562, depending on their associated promoter, identified using g:Profiler.

**Fig. S3.**

qPCR analysis of *PLEKHA4*, *PPP1R15A*, and *GYS1* expression in wild type and 4 ΔNUCB1 mutants in K562 cells before and after HS. Values represent the relative expression of the samples normalized to the housekeeping gene *GAPDH* and compared to the unstressed wild type. *P* values were calculated by a two-sided Student’s t-test, \*\*\**P* < 0.001, \*\**P* < 0.01, \**P* < 0.1.

**Fig. S4.**

(A) Screenshot of the GBP cluster by TNFα stimulation from the “Kusnadi” dataset. The genomic tracks show the ChIP-seq signal of SREBP after stimulation. The fold-change of induction is indicated below the name and orientation of the genes (Epromoter: green, co-induced genes: yellow). Note that all five *GBP* genes are induced but all excepted *GBP4* binds SREBP.

(B) (top panel) Mean normalized counts for *P2RX7/P2rx7* and *OASL/Oasl1/Oasl2* in unstimulated and IFNγ-stimulated human (blue) or mouse (green) macrophages. Data were retrieved from the “Langlais” dataset and from (*82*). (bottom panel) Schematic representation of the locus in the human (left) and mouse (right).

## References

1. B. Plosky, Stress. Mol Cell 84, 1 (2024).

2. J. C. Pessa, J. Joutsen, L. Sistonen, Transcriptional reprogramming at the intersection of the heat shock response and proteostasis. Molecular Cell 84, 80–93 (2024).

3. A. Vihervaara, F. M. Duarte, J. T. Lis, Molecular mechanisms driving transcriptional stress responses. Nature Reviews Genetics 19, 385–397 (2018).

4. L. Galluzzi, T. Yamazaki, G. Kroemer, Linking cellular stress responses to systemic homeostasis. Nature Reviews Molecular Cell Biology 19, 731–745 (2018).

5. A. T. Ghanbarian, L. D. Hurst, Neighboring Genes Show Correlated Evolution in Gene Expression. Mol Biol Evol 32, 1748–1766 (2015).

6. M. Ebisuya, T. Yamamoto, M. Nakajima, E. Nishida, Ripples from neighbouring transcription. Nat Cell Biol 10, 1106–1113 (2008).

7. A. Feuerborn, P. R. Cook, Why the activity of a gene depends on its neighbors. Trends Genet 31, 483–490 (2015).

8. W. Siwek, S. S. H. Tehrani, J. F. Mata, L. E. T. Jansen, Activation of Clustered IFNγ Target Genes Drives Cohesin-Controlled Transcriptional Memory. Molecular Cell 80, 396–409.e396 (2020).

9. D. M. Ribeiro, C. Ziyani, O. Delaneau, Shared regulation and functional relevance of local gene co-expression revealed by single cell analysis. Commun Biol 5, 876 (2022).

10. M. A. Zabidi et al., Enhancer–core-promoter specificity separates developmental and housekeeping gene regulation. Nature 518, 556–559 (2015).

11 . C. D. Arnold et al., Genome-wide quantitative enhancer activity maps identified by STARR-seq. Science 339, 1074–1077 (2013).

12. M. Corrales et al., Clustering of Drosophila housekeeping promoters facilitates their expression. Genome Res 27, 1153–1161 (2017).

13. T. A. Nguyen et al., High-throughput functional comparison of promoter and enhancer activities. Genome Research 26, 1023–1033 (2016).

14. J. M. Engreitz et al., Local regulation of gene expression by lncRNA promoters, transcription and splicing. Nature 539, 452–455 (2016).

15. L. T. M. Dao et al., Genome-wide characterization of mammalian promoters with distal enhancer functions. Nat Genet 49, 1073–1081 (2017).

16. Y. Diao et al., A tiling-deletion-based genetic screen for cis-regulatory element identification in mammalian cells. Nature Methods 14, 629–635 (2017).

17. N. Rajagopal et al., High-throughput mapping of regulatory DNA. Nat Biotechnol 34, 167–174 (2016).

18. D. Santiago-Algarra et al., Epromoters function as a hub to recruit key transcription factors required for the inflammatory response. Nature communications 12, 6660 (2021).

19. J. Malfait, J. Wan, S. Spicuglia, Epromoters are new players in the regulatory landscape with potential pleiotropic roles. Bioessays 45, e2300012 (2023).

20. F. Muerdter et al., Resolving systematic errors in widely used enhancer activity assays in human cells. Nature Methods 15, 141–149 (2018).

21. D. Wang et al., Comprehensive functional genomic resource and integrative model for the human brain. Science 362, eaat8464 (2018).

22. J. Mitchelmore, N. F. Grinberg, C. Wallace, M. Spivakov, Functional effects of variation in transcription factor binding highlight long-range gene regulation by epromoters. Nucleic Acids Research 48, 2866–2879 (2020).

23. I. Jung et al., A compendium of promoter-centered long-range chromatin interactions in the human genome. Nat Genet 51, 1442–1449 (2019).

24. M. Saint Just Ribeiro, P. Tripathi, B. Namjou, J. B. Harley, I. Chepelev, Haplotype-specific chromatin looping reveals genetic interactions of regulatory regions modulating gene expression in 8p23.1. Frontiers in genetics 13, 1008582 (2022).

25. L. T. M. Dao, S. Spicuglia, Transcriptional regulation by promoters with enhancer function. Transcription, (2018).

26. M. Dejosez et al., Regulatory architecture of housekeeping genes is driven by promoter assemblies. Cell Reports 42, 112505 (2023).

27. P. R. Cook, D. Marenduzzo, Transcription-driven genome organization: a model for chromosome structure and the regulation of gene expression tested through simulations. Nucleic Acids Research 46, 9895–9906 (2018).

28. A. Medina-Rivera, D. Santiago-Algarra, D. Puthier, S. Spicuglia, Widespread Enhancer Activity from Core Promoters. Trends in biochemical sciences 43, 452–468 (2018).

29. W. Schaffner, Enhancers, enhancers – from their discovery to today’s universe of transcription enhancers. Biological chemistry 396, 311–327 (2015).

30. A. Vihervaara et al., Transcriptional response to stress is pre-wired by promoter and enhancer architecture. Nature communications 8, 255 (2017).

31. C. Esnault et al., Rho-actin signaling to the MRTF coactivators dominates the immediate transcriptional response to serum in fibroblasts. Genes Dev 28, 943–958 (2014).

32. J. M. Ramos Pittol, A. Oruba, G. Mittler, S. Saccani, D. van Essen, Zbtb7a is a transducer for the control of promoter accessibility by NF-kappa B and multiple other transcription factors. PLOS Biology 16, e2004526 (2018).

33. S. C. Biddie et al., Transcription Factor AP1 Potentiates Chromatin Accessibility and Glucocorticoid Receptor Binding. Molecular Cell 43, 145–155 (2011).

34. V. Piccolo et al., Opposing macrophage polarization programs show extensive epigenomic and transcriptional cross-talk. Nat Immunol 18, 530–540 (2017).

35. A. Mancino et al., A dual cis-regulatory code links IRF8 to constitutive and inducible gene expression in macrophages. Genes Dev 29, 394–408 (2015).

36. D. B. Mahat, H. H. Salamanca, F. M. Duarte, C. G. Danko, J. T. Lis, Mammalian Heat Shock Response and Mechanisms Underlying Its Genome-wide Transcriptional Regulation. Mol Cell 62, 63–78 (2016).

37. X. Lyu, M. J. Rowley, V. G. Corces, Architectural Proteins and Pluripotency Factors Cooperate to Orchestrate the Transcriptional Response of hESCs to Temperature Stress. Molecular Cell 71, 940–955.e947 (2018).

38. S. F. Schmidt et al., Acute TNF-induced repression of cell identity genes is mediated by NFkappaB-directed redistribution of cofactors from super-enhancers. Genome Res 25, 1281–1294 (2015).

39. F. Gualdrini et al., SRF Co-factors Control the Balance between Cell Proliferation and Contractility. Mol Cell 64, 1048–1061 (2016).

40. J. D. Brown et al., NF-κB directs dynamic super enhancer formation in inflammation and atherogenesis. Mol Cell 56, 219–231 (2014).

41. M. D. Cardamone et al., Mitochondrial Retrograde Signaling in Mammals Is Mediated by the Transcriptional Cofactor GPS2 via Direct Mitochondria-to-Nucleus Translocation. Mol Cell 69, 757–772.e757 (2018).

42. C. Camps et al., Integrated analysis of microRNA and mRNA expression and association with HIF binding reveals the complexity of microRNA expression regulation under hypoxia. Mol Cancer 13, 28 (2014).

43. F. Jin et al., A high-resolution map of the three-dimensional chromatin interactome in human cells. Nature 503, 290–294 (2013).

44. A. W. Jubb, R. S. Young, D. A. Hume, W. A. Bickmore, Enhancer Turnover Is Associated with a Divergent Transcriptional Response to Glucocorticoid in Mouse and Human Macrophages. J Immunol 196, 813–822 (2016).

45. N. T. Hogan et al., Transcriptional networks specifying homeostatic and inflammatory programs of gene expression in human aortic endothelial cells. Elife 6, (2017).

46. M. L. Hancock et al., Insulin Receptor Associates with Promoters Genome-wide and Regulates Gene Expression. Cell 177, 722–736.e722 (2019).

47. H. L. Franco, A. Nagari, W. L. Kraus, TNFalpha signaling exposes latent estrogen receptor binding sites to alter the breast cancer cell transcriptome. Mol Cell 58, 21–34 (2015).

48. A. Kusnadi et al., The Cytokine TNF Promotes Transcription Factor SREBP Activity and Binding to Inflammatory Genes to Activate Macrophages and Limit Tissue Repair. Immunity 51, 241–257.e249 (2019).

49. R. Ferrari et al., TFIIIC Binding to Alu Elements Controls Gene Expression via Chromatin Looping and Histone Acetylation. Mol Cell 77, 475–487.e411 (2020).

50. D. Langlais, L. B. Barreiro, P. Gros, The macrophage IRF8/IRF1 regulome is required for protection against infections and is associated with chronic inflammation. J Exp Med 213, 585–603 (2016).

51. K. A. Lo et al., Analysis of in vitro insulin-resistance models and their physiological relevance to in vivo diet-induced adipose insulin resistance. Cell Rep 5, 259–270 (2013).

52. J. R. Porter et al., Global Inhibition with Specific Activation: How p53 and MYC Redistribute the Transcriptome in the DNA Double-Strand Break Response. Mol Cell 67, 1013–1025.e1019 (2017).

53. S. H. Park et al., Type I interferons and the cytokine TNF cooperatively reprogram the macrophage epigenome to promote inflammatory activation. Nat Immunol 18, 1104–1116 (2017).

54. D. H. Phanstiel et al., Static and Dynamic DNA Loops form AP-1-Bound Activation Hubs during Macrophage Development. Mol Cell 67, 1037–1048.e1036 (2017).

55. T. Vierbuchen et al., AP-1 Transcription Factors and the BAF Complex Mediate Signal-Dependent Enhancer Selection. Mol Cell 68, 1067–1082.e1012 (2017).

56. B. E. Bernstein et al., The NIH Roadmap Epigenomics Mapping Consortium. Nat Biotechnol 28, 1045–1048 (2010).

57. M. Gasperini et al., A Genome-wide Framework for Mapping Gene Regulation via Cellular Genetic Screens. Cell 176, 377–390 e319 (2019).

58. D. M. Ribeiro et al., The molecular basis, genetic control and pleiotropic effects of local gene co-expression. Nature communications 12, 4842 (2021).

59. M. Mineo et al., Tumor Interferon Signaling Is Regulated by a lncRNA INCR1 Transcribed from the PD-L1 Locus. Mol Cell 78, 1207–1223 e1208 (2020).

60. C. Sun, R. Mezzadra, T. N. Schumacher, Regulation and Function of the PD-L1 Checkpoint. Immunity 48, 434–452 (2018).

61. B. M. Javierre et al., Lineage-Specific Genome Architecture Links Enhancers and Non-coding Disease Variants to Target Gene Promoters. Cell 167, 1369–1384 e1319 (2016).

62. J. O. Onuh, H. Qiu, Serum response factor-cofactor interactions and their implications in disease. FEBS J 288, 3120–3134 (2021).

63. A. Field, K. Adelman, Evaluating Enhancer Function and Transcription. Annu Rev Biochem 89, 213–234 (2020).

64. R. Andersson, A. Sandelin, Determinants of enhancer and promoter activities of regulatory elements. Nat Rev Genet 21, 71–87 (2020).

65. R. R. Catarino, C. Neumayr, A. Stark, Promoting transcription over long distances. Nat Genet 49, 972–973 (2017).

66. T. Henriques et al., Widespread transcriptional pausing and elongation control at enhancers. Genes Dev 32, 26–41 (2018).

67. O. Mikhaylichenko et al., The degree of enhancer or promoter activity is reflected by the levels and directionality of eRNA transcription. Genes & Development 32, 42–57 (2018).

68. S. Rennie et al., Transcription start site analysis reveals widespread divergent transcription in D. melanogaster and core promoter-encoded enhancer activities. Nucleic Acids Res 46, 5455–5469 (2018).

69. L. J. Core et al., Analysis of nascent RNA identifies a unified architecture of initiation regions at mammalian promoters and enhancers. Nat Genet 46, 1311–1320 (2014).

70. S. Fanucchi, Y. Shibayama, S. Burd, M. S. Weinberg, M. M. Mhlanga, Chromosomal contact permits transcription between coregulated genes. Cell 155, 606–620 (2013).

71. S. Chowdhary, A. S. Kainth, S. Paracha, D. S. Gross, D. Pincus, Inducible transcriptional condensates drive 3D genome reorganization in the heat shock response. Mol Cell 82, 4386–4399.e4387 (2022).

72. P. Bhat, D. Honson, M. Guttman, Nuclear compartmentalization as a mechanism of quantitative control of gene expression. Nature reviews. Molecular cell biology 22, 653–670 (2021).

73. S. Cuartero et al., Control of inducible gene expression links cohesin to hematopoietic progenitor self-renewal and differentiation. Nat Immunol 19, 932–941 (2018).

74. A. E. Shaw et al., Fundamental properties of the mammalian innate immune system revealed by multispecies comparison of type I interferon responses. PLoS Biol 15, e2004086 (2017).

75. X. Wu, P. A. Sharp, Divergent transcription: a driving force for new gene origination? Cell 155, 990–996 (2013).

76. R. Andersson, Promoter or enhancer, what’s the difference? Deconstruction of established distinctions and presentation of a unifying model: Prospects & Overviews. BioEssays 37, 314–323 (2015).

77. C. Arenas-Mena, The origins of developmental gene regulation. Evol Dev 19, 96–107 (2017).

78. C. Xie et al., Hominoid-specific de novo protein-coding genes originating from long non-coding RNAs. PLoS genetics 8, e1002942 (2012).

79. F. N. Carelli, A. Liechti, J. Halbert, M. Warnefors, H. Kaessmann, Repurposing of promoters and enhancers during mammalian evolution. Nature communications 9, 4066 (2018).

80. P. Majic, J. L. Payne, Enhancers Facilitate the Birth of De Novo Genes and Gene Integration into Regulatory Networks. Molecular Biology and Evolution 37, 1165–1178 (2020).

81. S. E. Gilbertson et al., Topologically associating domains are disrupted by evolutionary genome rearrangements forming species-specific enhancer connections in mice and humans. Cell Reports 39, 110769 (2022).

82. K. Kang et al., Interferon-γ Represses M2 Gene Expression in Human Macrophages by Disassembling Enhancers Bound by the Transcription Factor MAF. Immunity 47, 235–250.e234 (2017).

83. M. G. Netea et al., A guiding map for inflammation. Nat Immunol 18, 826–831 (2017).

84. V. Chandra et al., Promoter-interacting expression quantitative trait loci are enriched for functional genetic variants. Nat Genet 53, 110–119 (2021).

85. V. Rusu et al., Type 2 Diabetes Variants Disrupt Function of SLC16A11 through Two Distinct Mechanisms. Cell 170, 199–212.e120 (2017).

86. I. A. Sergeeva et al., Identification of a regulatory domain controlling the *Nppa-Nppb* gene cluster during heart development and stress. Development, dev.132019 (2016).

87. N. Yagihara et al., Variants in the SCN5A Promoter Associated With Various Arrhythmia Phenotypes. J Am Heart Assoc 5, (2016).

88. S. Nisar et al., Identification of ATP2B4 Regulatory Element Containing Functional Genetic Variants Associated with Severe Malaria. International journal of molecular sciences 23, 4849 (2022).

89. J. T. Hua et al., Risk SNP-Mediated Promoter-Enhancer Switching Drives Prostate Cancer through lncRNA PCAT19. Cell 174, 564–575 e518 (2018).

90. P. Gao et al., Biology and Clinical Implications of the 19q13 Aggressive Prostate Cancer Susceptibility Locus. Cell 174, 576–589 e518 (2018).

91. J. Reimand et al., g:Profiler-a web server for functional interpretation of gene lists (2016 update). Nucleic Acids Res, (2016).

92. S. Sayols, rrvgo: a Bioconductor package for interpreting lists of Gene Ontology terms. MicroPubl Biol 2023, (2023).

93. D. Lin et al., Decoding the spatial chromatin organization and dynamic epigenetic landscapes of macrophage cells during differentiation and immune activation. Nature communications 13, 5857 (2022).

94. D. Li et al., WashU Epigenome Browser update 2022. Nucleic Acids Res 50, W774–W781 (2022).

