## Supplementary Figures for "Epromoters bind key stress-related transcription factors to regulate clusters of stress response genes"

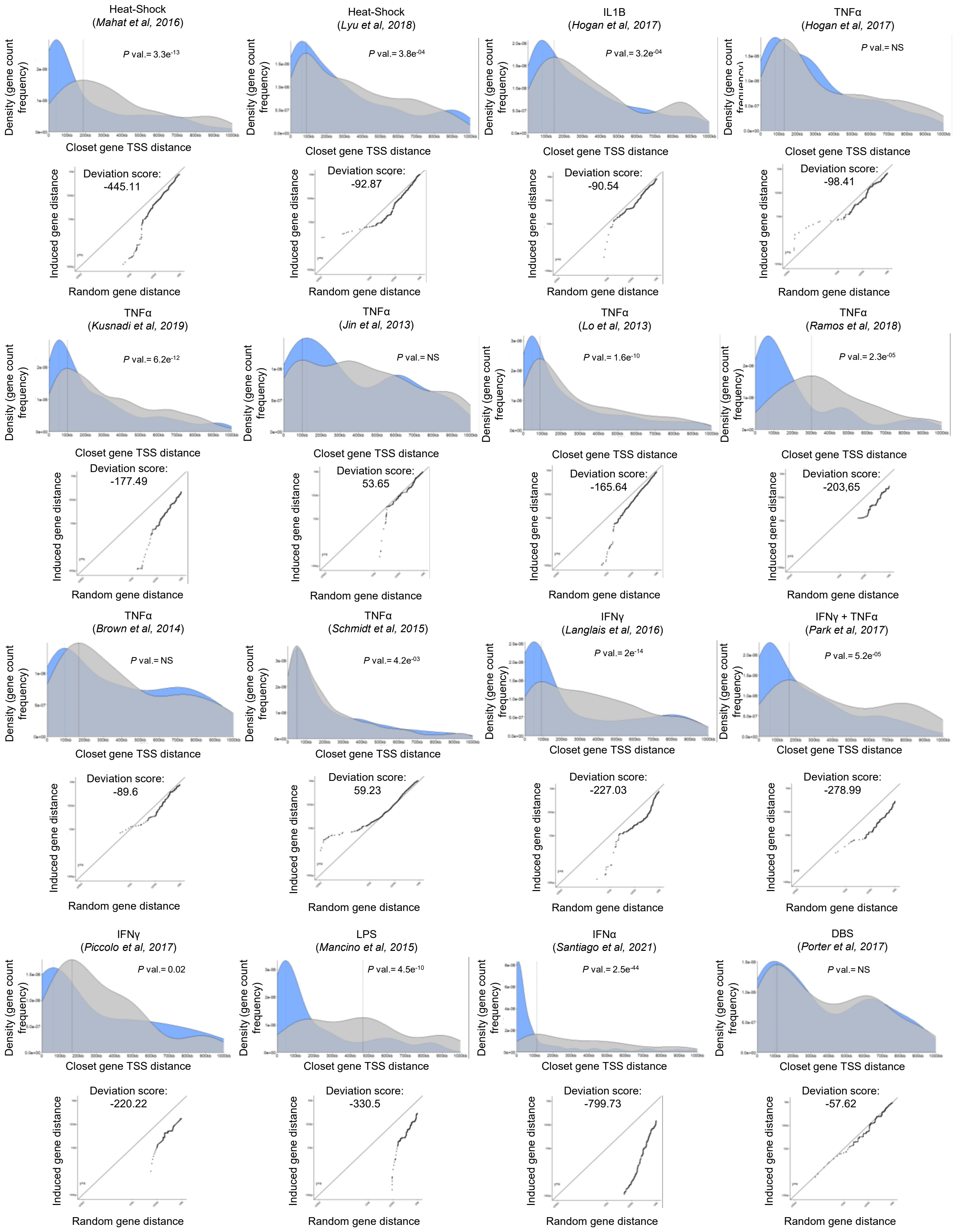


Supplementary Figure S1 (part 1)

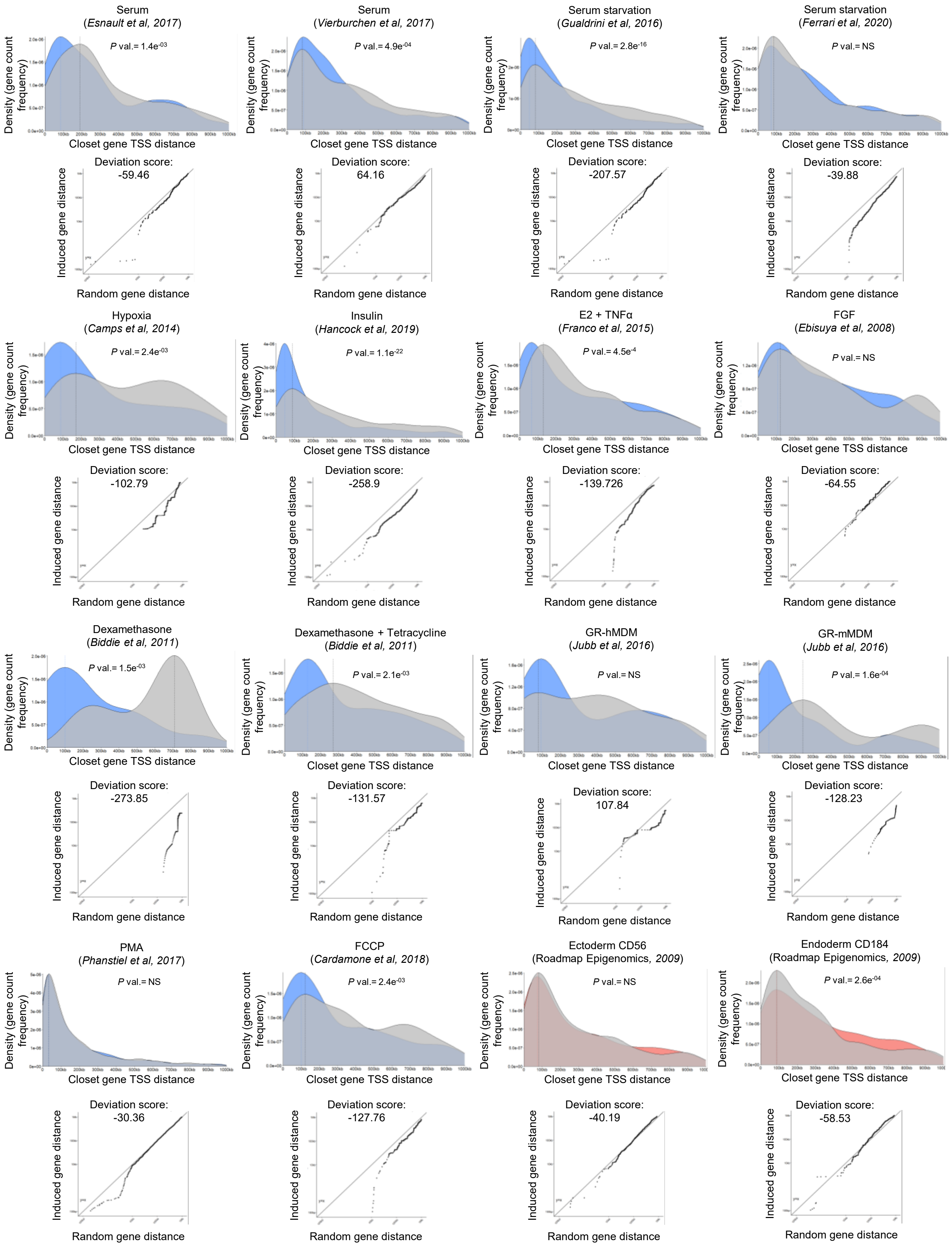


Supplementary Figure S1 (part 2)

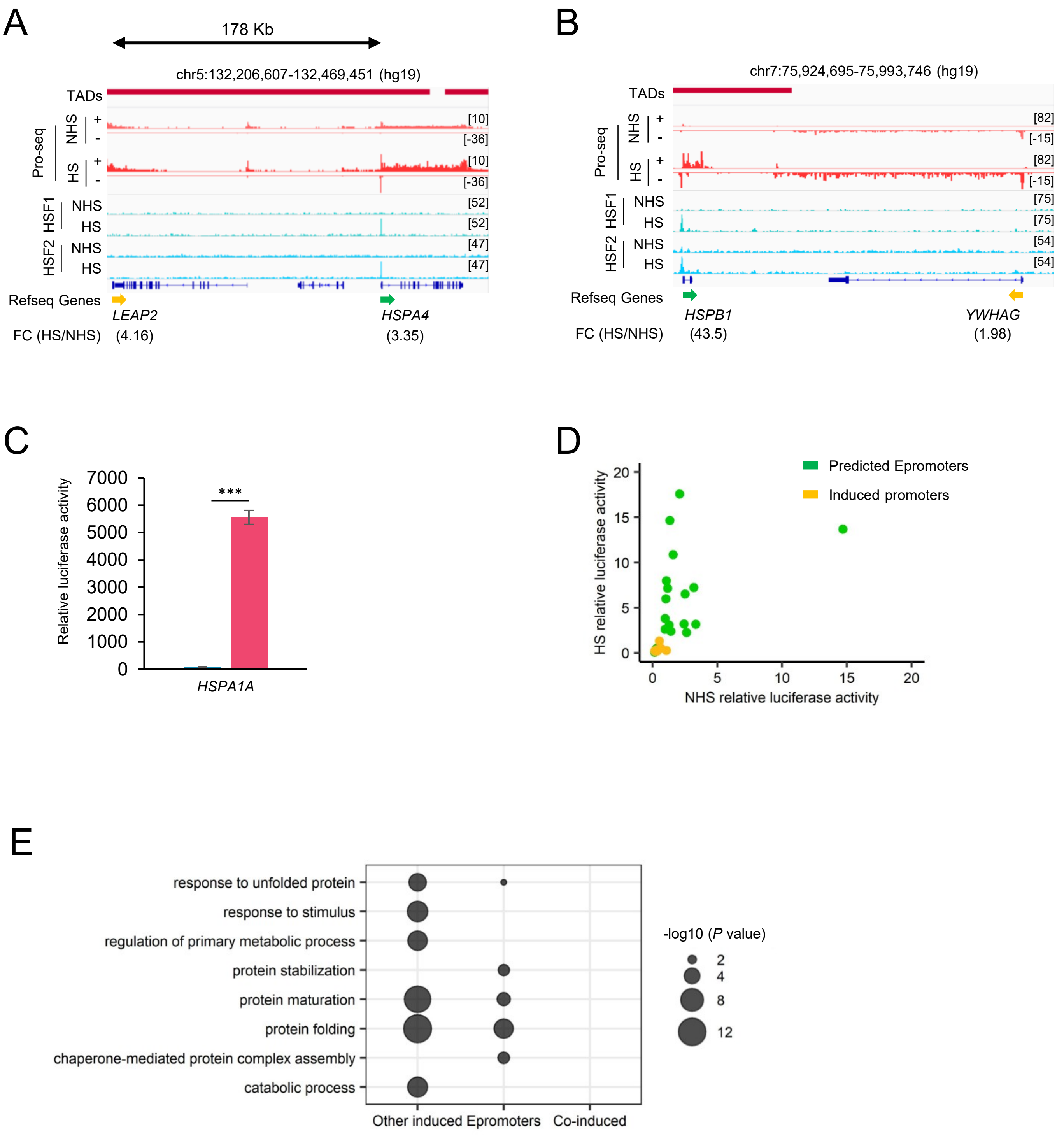


Supplementary Figure S2

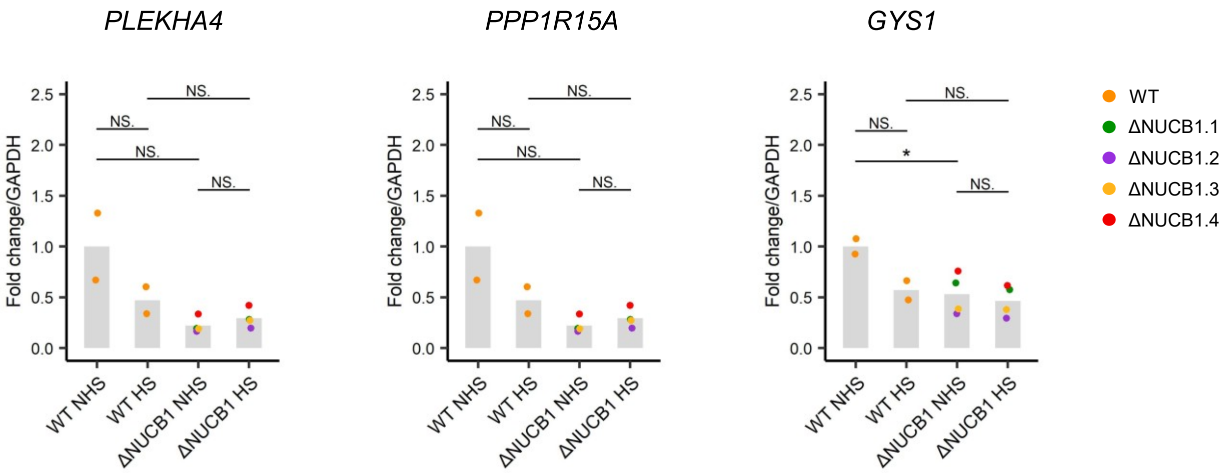


Supplementary Figure S3

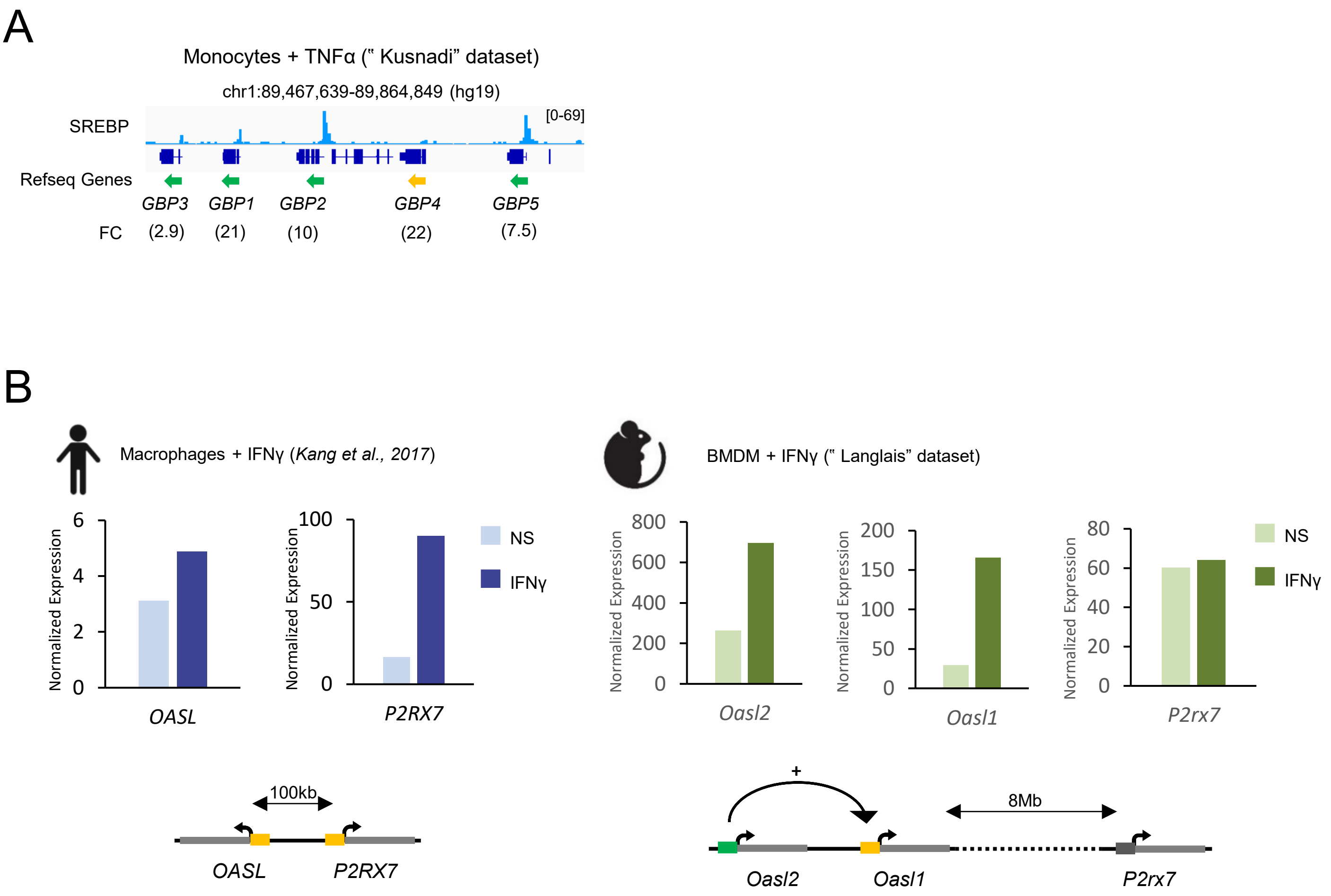


Supplementary Figure S4
